# A Failure Mode and Effect Analysis of plant metabolism reveals why cytosolic fumarase is required for temperature acclimation in Arabidopsis

**DOI:** 10.1101/2020.08.04.234591

**Authors:** Helena A. Herrmann, Pablo I. Calzadilla, Jean-Marc Schwartz, Giles N. Johnson

## Abstract

Plants acclimate their photosynthetic capacity in response to changing environmental conditions. In *Arabidopsis thaliana*, photosynthetic acclimation to cold requires the accumulation of the organic acid fumarate, catalysed by a cytosolic fumarase FUM2, however the role of this is currently unclear.

In this study, we use an integrated experimental and modelling approach to examine the role of FUM2 and fumarate across the physiological temperature range. Using physiological and biochemical analyses, we demonstrate that FUM2 is necessary for high as well as low temperature acclimation.

To understand the role of FUM2 activity, we have adapted a reliability engineering technique, Failure Mode and Effect Analysis (FMEA), to formalize a rigorous approach for ranking metabolites according to the potential risk that they pose to the metabolic system. FMEA identifies fumarate as a low-risk metabolite. Its precursor, malate, is shown to be high-risk and liable to cause system instability. We conclude that the role of cytosolic fumarase, FUM2, is to provide a fail-safe, maintaining system stability under changing environmental conditions.

We argue that FMEA is a technique which is not only useful in understanding plant metabolism, it can also be used to study reliability in other systems and aid the design of synthetic pathways.

## Introduction

Metabolism is the basis of all cellular. Metabolic systems must be optimized to maintain function, regardless of the environmental conditions experienced. Plants are exposed to environments which fluctuate over timescales varying from seconds to decades, with environmental parameters such as light and temperature directly impacting metabolism. As such, plants provide an excellent example in which to understand how metabolic systems can be optimized to tolerate environmental change. To understand such optimization, we can use approaches taking form system engineering.

In system processes, the quality of a function or product is referred to as its reliability. The quantity, on the other hand, is dependent on the system’s robustness (King and Jewett, 2010). As an analogy, we can consider a business process where the aim of the process is to produce as many high-quality products as possible, despite fluctuations in operating conditions. A reliable process ensures that the quality of the products is consistently high, whereas a robust process ensures that as many products as possible are being produced. If all low-quality products are discarded then, to make a profit, the system must first be reliable and then robust (Bakera *et al*., 2008; King and Jewett, 2010). This is comparable to plants optimizing seed production (Sadras 2007). Only viable seeds will germinate; thus, features of reliability, which ensure that seeds germinate, are of primary concern. Features of robustness, which ensure that as many seeds as possible are produced, are secondary. In fact, in the model plant species *Arabidopsis thaliana* the trade-off between seed size and number is minimal (Gnan *et al*., 2014), suggesting that these traits may be selected for independently.

This distinction between robustness and reliability is considered crucial in many disciplines, including systems engineering and product development (Yassine, 2007; King and Jewett, 2010). With regards to metabolism, the two concepts are rarely distinguished, and the reliability of metabolic networks has not been formally quantified. However, reliability engineering frameworks exist which can be applied to metabolic networks.

Failure mode and effect analysis (FMEA) is a framework that is commonly applied to study the reliability of a product of a process, and is considered to be one of the most effective ways of conducting a failure analysis (i.e. collecting data on why a system might fail). FMEA breaks the system down into individual components or single processes and quantifies the risk that each of them poses to the system (Stamatis, 1995; Rausanda and Øienb, 1996). The risk of each component (or process) is assessed by studying the potential causes of failure and effects that these would have on the system. To quantify this risk, each component is assigned a probability of failure (*P*) and a severity score in the event of failure (*S*). The risk (*R*) of each component is then calculated such that *R = P × S*. Components which are essential for the system to function are considered high-risk. A well-engineered system has control mechanisms to regulate the functioning of the high-risk components, to minimize the impact of failures. Such control mechanisms can be inherent features of the system (e.g. the closing of one valve automatically opens another) or can be sensed and regulated (e.g. a sensor perceives that one valve is closed and opens another).

The benefit of the FMEA framework is that it can be adapted to conduct a failure analysis of any system, making it also suitable for studies of metabolic systems. We can think of individual metabolites as the components that constitute a system. The metabolic system may represent the whole of metabolism or a subset of metabolites, depending on the processes of interest. The components (i.e. the metabolites) need to be present within specific concentration ranges, or the system will not function correctly. Failure of the system might mean a failure to produce certain products at an appropriate rate (e.g. a failure to synthesise sufficient ATP to support cell maintenance) or the accumulation of metabolites to toxic concentrations. We would expect the concentration of high-risk metabolites to be tightly regulated to avoid systems failures. The probability of failure of an individual metabolite, i.e. the probability that its concentration falls outside an acceptable range, could be considered a function of upstream metabolites and the severity of failure as a function of downstream metabolites. Thus, the risk score for each metabolite arises from the underlying metabolic network structure, the highest risk metabolites being those which are heavily dependent on the production of other metabolites whilst at the same time supporting the production of many others.

Reliability engineering has potential in trying to understand how metabolic systems respond to environmental stress. For example, it is predicted that, due to anthropogenic climate change, plants will increasingly be exposed to extreme weather events, including both high and low temperature periods (Changnon *et al*., 2000; Trenberth, 2012). These extreme conditions will affect crops yield, threatening food security for a growing world population (Lesk *et al*., 2016; Powell and Reinhard, 2016). Thus, the identification of traits which determine the reliability of key acclimation processes becomes a priority. In this context, plant responses to single temperature stresses have been well characterized (e.g. Hikosaka *et al*., 2006; Yamori *et al*., 2014; Ding *et al*., 2020). A few studies have also compared the effects of warm and cold temperature stresses on Arabidopsis leaves, showing that the two processes share common features (Kaplan *et al*., 2004; Zhang *et al*., 2019). Nevertheless, how metabolism adjusts across the entire physiological temperature range of a plant is still largely unknown.

When exposed to short term environmental changes, plants can *regulate* enzymes to ensure metabolic function is maintained. Over longer periods, plants *acclimate* to their environment, altering the concentration of different enzymes to match the prevailing conditions (Webber, 1994; Stitt and Hurry, 2002). For example, when a plant is transferred from warm to cold conditions, enzyme activities are slowed, reducing metabolic activity. This would be reflected, for example, in a reduction in the light-saturated rate of photosynthesis. Following acclimation, plants transferred to the cold typically increase the concentration of key enzymes, allowing them to achieve a similar rate of photosynthesis at low temperature to that previously seen in the warm (Hurry *et al*., 2000; Strand *et al*., 2003; Dyson *et al*., 2016). This results in an increase in the photosynthetic capacity (*P*_*max*_) (Athanasiou *et al*., 2010; Dyson *et al*., 2016). Conversely, in response to increased temperature, plants decrease their *P*_*max*_, allowing them to reallocate resources to other processes (Herrmann *et al*., 2019b).

During the day, plants fix atmospheric carbon through photosynthesis. A large proportion of that fixed carbon is accumulated in the leaf during the photoperiod. Overnight this is then remobilised to support growth and cell maintenance. Under different environmental conditions, plants vary the form in which photoassimilates are stored. Arabidopsis stores carbon primarily in the form of the polysaccharide starch and the organic acids fumarate and malate (Chia *et al*., 2000; Smith and Stitt, 2007; Zell *et al*., 2010; Pracharoenwattana *et al*., 2010). Starch accumulation and degradation have been studied extensively as this is the major non-structural carbohydrate in plants (Smith and Stitt, 2007; Streb and Zeeman, 2012). Less is known about malate and fumarate. Both metabolites are intermediates of the tricarboxylic acid cycle in the mitochondria; however, it has also been shown that they accumulate in the vacuole diurnally (Kovermann *et al*., 2007; Fernie and Martinoia, 2009). In the cytosol, malate is made from or converted to oxaloacetate, citrate, isocitrate, cis-aconitate, fumarate, α-ketaglutarate, glutamic acid and pyruvate (Arnold and Nikoloski, 2014). It provides carbon skeletons for various biosynthetic pathways, including amino acid synthesis (Zell *et al*., 2010). In the cytosol, fumarate is synthesised by a cytosolic isoform of the enzyme fumarase, FUM2 (Pracharoenwattana *et al*., 2010). It is made from malate during the day and is assumed to be converted back to malate during the night (Pracharoenwattana *et al*., 2010; Zell *et al*., 2010; Dyson *et al*., 2016). Fumarate accumulation in the cytosol has previously been linked to growth on high nitrogen (Pracharoenwattana *et al*., 2010) and photosynthetic acclimation to cold (Dyson *et al*., 2016); however, its role in these processes is unclear (Chia *et al*., 2000).

The *fum2* Arabidopsis mutant is a cytosolic fumarase knockout (Pracharoenwattana *et al*., 2010). Previously, we showed that fumarate and malate accumulation increase in response to cold, and that the *fum2* knockout mutants, which do not accumulate fumarate, are unable to acclimate photosynthetic capacity in response to low temperature (Dyson *et al*., 2016). The mutant accumulates higher levels of both malate and starch under cold stress. It has also been shown that plants which accumulate more fumarate accumulate less starch (Chia *et al*., 2000; Pracharoenwattana *et al*., 2010; Dyson *et al*., 2016) and that fumarate accumulation is proportional to biomass production at low temperature (Scott *et al*., 2014). Therefore, fumarate accumulation participates in the cold acclimation response, although its specific role in this remains to be elucidated.

In this study, we demonstrate that FUM2 activity is essential not only for low temperature acclimation but also for high temperature responses. Using three genotypes, with varying capacities to accumulate fumarate, we have examined acclimation across the physiological temperature range. We have measured photosynthetic and metabolic responses of the Col-0 wild-type, which accumulates high levels of fumarate, a *fum2* knockout, which does not accumulate fumarate (Pracharoenwattana *et al*., 2010), and the C24 wild-type, which accumulates intermediate levels of fumarate compared to Col-0 (Riewe *et al*., 2016). To understand their responses to temperature treatment, we combine the novel application of FMEA with a more widely used kinetic modelling approach (Saa and Nielsen, 2017; Herrmann et al., 2019a), identifying cytosolic malate as a high-risk component and its metabolic neighbour, cytosolic fumarate, as a low-risk one. We propose that fumarate synthesis acts as a fail-safe, maintaining malate concentrations within safe limits and ensuring that metabolic functions continue across a wide range of conditions. We discuss that FMEA is a valuable approach to understanding metabolic system with possible widespread applications in biotechnology and systems biology.

## Materials and Methods

### Plant growth and tissue preparation

*A. thaliana* Col-0, *fum2*.*2*, and C24 genotypes were grown in a multi-purpose peat-based compost in 3-inch pots with 8-hour photoperiods at 20 °C and under a 100 μmol m^-2^ s^-1^ irradiance, as previously described (Dyson *et al*., 2016). Light was provided by warm white LEDs (colour temperature 2800-3200 K). Adult plants (8-9 weeks) were then either kept at 20 °C for up to another week (control treatment) or transferred to 5, 10, 15, 25, or 30 °C, under identical irradiance and photoperiod conditions. Fully expanded leaves were collected at the beginning and at the end of the photoperiod on the 1^st^ and 7^th^ day of the different treatments, flash-frozen in liquid nitrogen, freeze-dried and stored at −20 °C for metabolite assays. Leaves were also collected without freezing and their fresh and dry weights per unit area recorded.

### Gas exchange

Photosynthesis and respiration were measured under growth conditions in plant growth cabinets, using a CIRAS 1 infra-red gas analyser fitted with a standard broad leaf chamber (PP systems, Amesbury, USA), as previously described (Dyson *et al*., 2016). Measurements of gas exchange under ambient growth conditions were taken on the 1^st^ and 7^th^ day for the cold (5 °C) and warm (30 °C) treatment, and under control conditions. The maximum capacity for photosynthesis (*P*_*max*_) was estimated at 20 °C and under 2000 ppm CO_2_ and 2000 µmol m^-2^ s^-1^ light, as described previously (Herrmann *et al*., 2019b). All measurements were taken between the 5-7^th^ hour of the photoperiod and on a minimum of 4 biological replicates.

### Metabolite assays

Fumarate, malate and starch concentrations were estimated as previously described (Dyson *et al*., 2016). Averages and standard errors of 3-4 replicates were calculated, and an analysis of variance (ANOVA) followed by a Turkey’s test was performed in R, with a confidence level of 0.95. Where measurements were subtracted from one another, the uncertainty (*μ*) was calculated such that 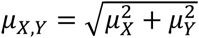. Measurements where the uncertainty ranges do not overlap are considered to be significantly different.

### Failure Mode and Effect Analysis (FMEA)

Metabolism can be represented in the form of a metabolic network (or graph), such that metabolites (nodes) are connected to one another (i.e. share an edge) if there exists a metabolic reaction that converts one metabolite to the other (Jeong *et al*., 2000). If the graph is undirected, all reactions are considered to be reversible. If the graph is directed, edges connected nodes only in the direction that the reaction is presumed to proceed. An existing genome-scale metabolic model of Arabidopsis plant leaf metabolism, constructed by Arnold and Nikoloski (2014) and updated by Herrmann *et al*. (2019c) was converted to a directed metabolite-metabolite graph and analysed using the *networkx* package (Version 2.2) in Python (Version 3.6.9). Reversible reactions were accounted for by applying two separate edges between the respective metabolites, one in either direction. All non-carbon compounds, small molecules and cofactors were removed from the graph. The model is leaf specific and is compartmentalized to include the chloroplast, mitochondrion, cytosol and peroxisomes.

We applied the FMEA framework to primary carbon metabolism. We considered metabolites as the components of the metabolic system. We calculated the risk of failure for each component considering, for instance, failure to be that the concentration of a particular metabolite falls outside a concentration range required to sustain metabolic activity. The FMEA analysis does not presume what the required concentration of a metabolite should be at a given condition and can therefore be considered an unbiased approach (Lewis *et al*., 2012).

The probability of failure for a given metabolite *M* downstream of ribulose-1,5-bisphosphate carboxylase/oxygenase (Rubisco) was calculated as:

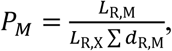

where *L*_*R,M*_ is equal to the shortest path length from Rubisco to metabolite *M, L*_*R,X*_ is the longest of all the shortest paths to metabolites in the network, and ∑ *d*_R,M_ is the number of alternative paths to metabolite M having the shortest length. *L*_*R,X*_ is a normalization constant and is equal to 14 in all calculation of *P*_*M*_. Overall, *P*_*M*_ considers the probability of failure of *M* as a calculation of the number of upstream reactions and metabolites that are required for the production of *M*. The severity of failure for each metabolite (*S*_*M*_) was calculated such that:

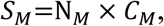

where *N*_*M*_ is the degree of metabolite *M* (i.e. number of direct neighbours) and *C*_*M*_ is the normalized betweenness centrality of *M*. The number of neighbours of *M* indicates the number of reactions that would fail if *M* was not present in the system. Betweenness centrality is defined as the normalized sum of the total number of shortest paths that pass through a node and can thus be considered a proxy for how important that metabolite is in the production of other metabolites. *S*_*M*_ calculates a severity for *M* based of the number of reactions and down-stream metabolites that cannot occur in the absence of *M*.

The risk factor (*R*_*M*_) was calculated for each metabolite such that

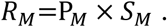

The risk factor takes into consideration how heavily the production of *M* is dependent on other metabolites and other reactions in the system. It also considers how many other metabolites and reactions in the system are dependent on the production of *M. R*_*M*_ therefore considers how important *M* is to the overall system functionality.

Although we could equally well calculate the risk factor of individual reactions in a metabolic system, we have here chosen to consider metabolites as the individual components of the system. This is because metabolites are the fundamental building blocks of the metabolic system, whereas reactions are the means to producing these building blocks. For example, it is possible for one reaction to fail but for all metabolites in the system to still be produced as required.

### Kinetic modelling

Kinetic modelling, unlike FMEA, can be used to describe changes in metabolite concentrations overtime. A set of mass-balanced reactions can be described by ordinary differential equations which capture the system dynamics (Saa and Nielsen, 2017; Herrmann et al., 2019a). Here, we used kinetic modelling to assess how changes in flux correspond to changes in metabolite concentrations. Kinetic modelling is not practical with large networks so, rather than using a genome-scale model we constructed a minimal model based on a subset of reactions (Fig. 1; Table S1). For simplicity we did not consider compartmentalization in the kinetic model. Kinetic reactions were set up in COPASI (Version 4.27.217). Rates of photosynthesis and respiration (Fig. 2b,c; Fig. S1) were set as independent variables for Col-0, *fum2*, and C24 plants, after being converted to μmol CO_2_ (gDW)^-1^ s^-1^ (Fig. S2). We initially parametrized the model to fit the measured average diurnal carbon fluxes to starch, malate, and fumarate for control conditions (Fig. 3) using simple mass action kinetics (Abegg, 1899). All other metabolites were assumed not to accumulate in the leaf and were constrained to concentrations between 0-0.0005 μmol CO_2_ (gDW)^1^ s ^-1^. The rate of carbon export (Fig.1) was allowed to adjust freely, accounting for any remaining carbon. Using the Hooke and Jeeves (1961) parameter estimation algorithm, with an interation limit of 10 000, a tolerance of 10^−8^ and a rho of 0.2, we found a model solution for which estimated concentration values fell within the uncertainty ranges of the experimentally measured values. We used an Arrhenius constant to capture temperature-dependence and allowed the effective *Q*_*10*_ to vary between 1.0-3.0 in order to fit the model to the measured starch, malate, and fumarate concentrations at *T* = 5 °C and *T* = 30 °C (Fig. 3). The Arrhenius constant describes an exponential increase of reaction rates with temperature (Arrhenius, 1889); it is most commonly reported for a 10 °C change in temperature, known as *Q*_*10*_, with a *Q*_*10*_ = 2 being typical for enzyme catalysed reactions (Elias, 2014). Without implementing any regulatory mechanisms, we were able to find a solution for which the concentration values estimated by the model fell within the uncertainty ranges of the experimentally measured values (Fig. S3). It is important to note that the effective Q_10_, as represented in our model, quantifies a possible temperature sensitivity of a reaction rather than the intrinsic properties of enzymes.

**Figure 1:**
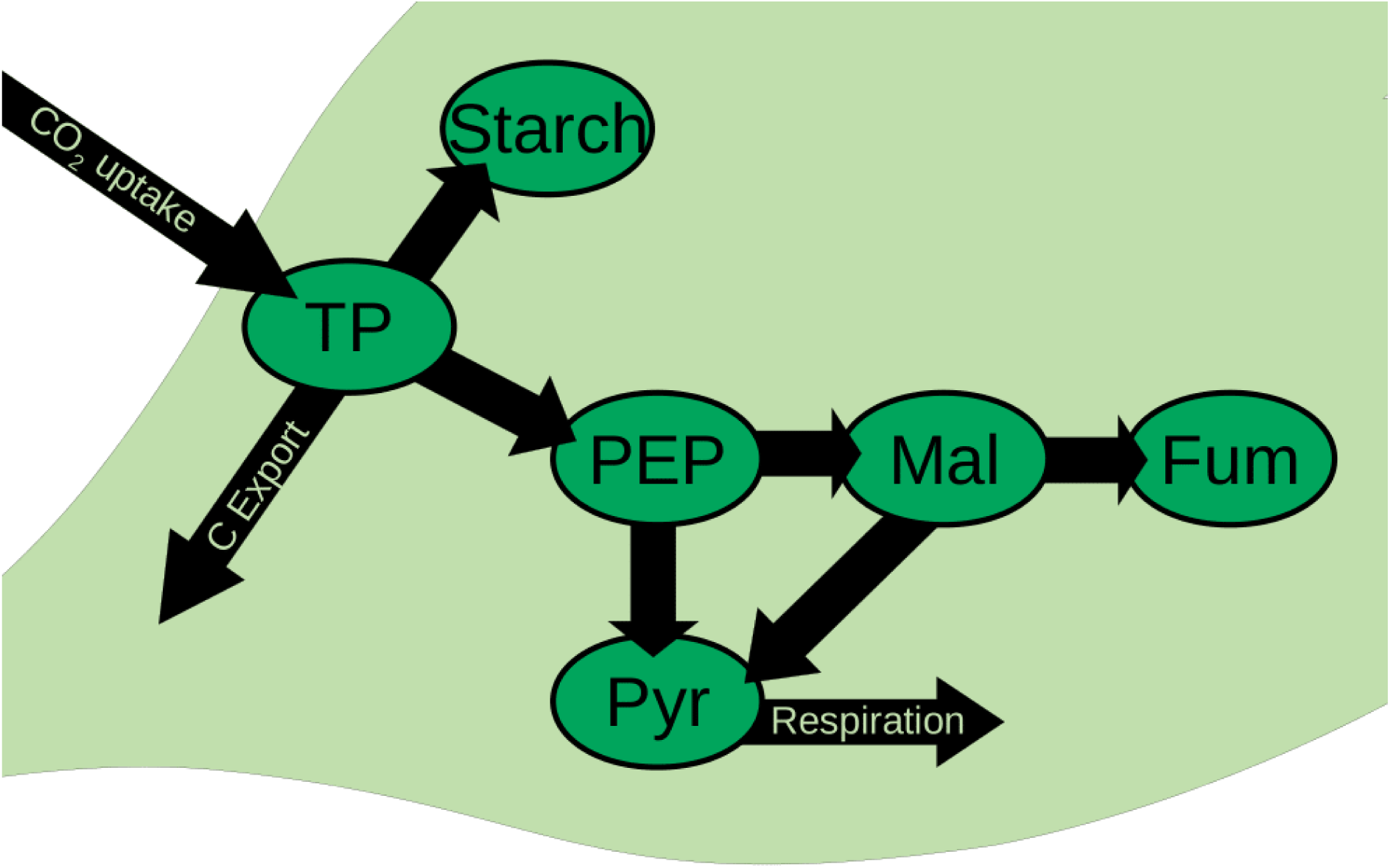
Illustration of the reactions used to set up the kinetic model. Reactions include the metabolites triose phosphate (TP), phosphoenolpyruvate (PEP), pyruvate (Pyr), malate (Mal), fumarate (Fum) and starch. The rate of photosynthesis (CO_2_ uptake) and respiration were constrained using experimentally measured values as outline in the Material and Methods. Model parameters were fitted to match the observed malate, fumarate and starch concentrations over an 8 h photoperiod on the first day of temperature treatment. All other metabolites were not allowed to accumulate in the leaf and all remaining carbon was assumed to be exported from the leaf (C export).

**Figure 2:**
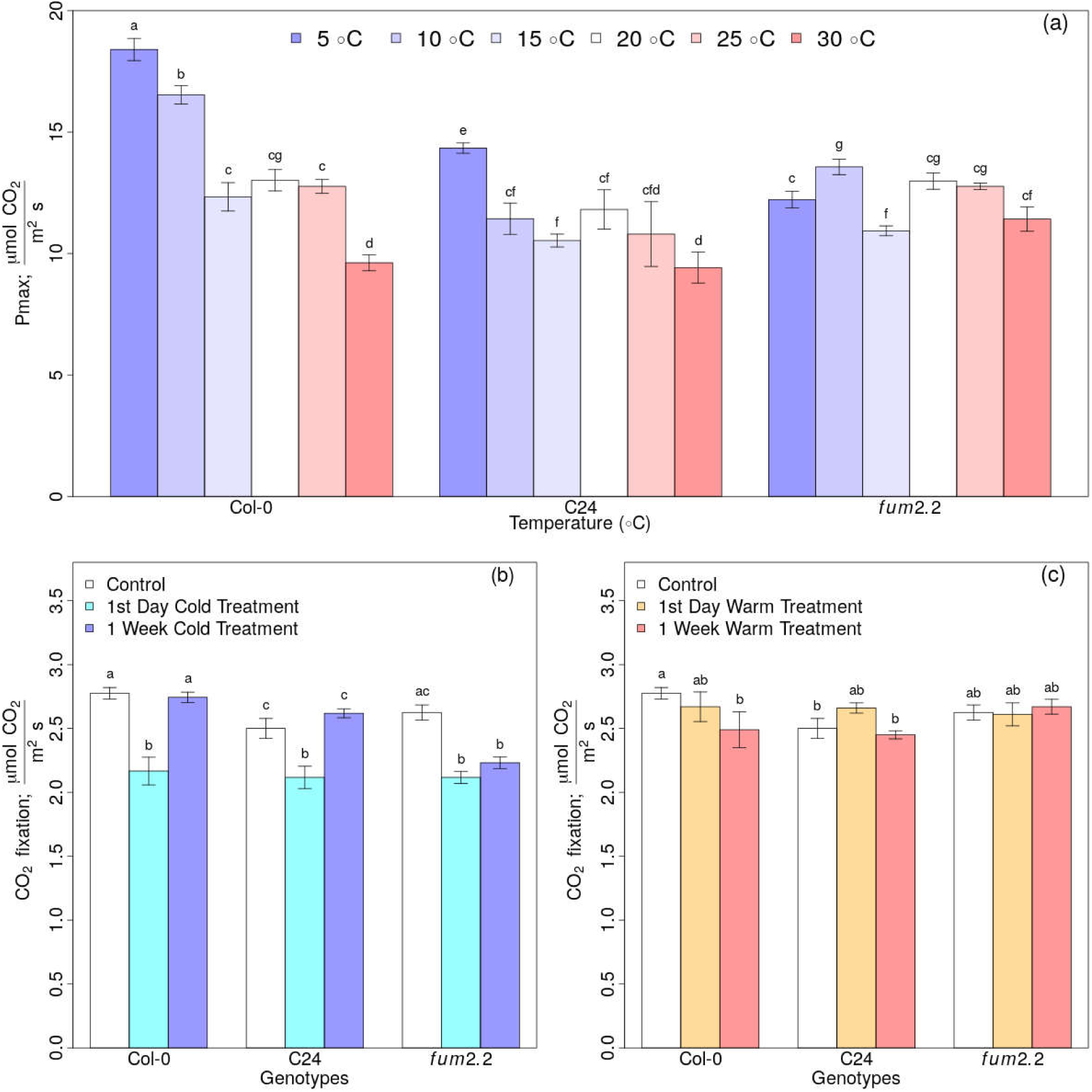
Effect of temperature on carbon assimilation in Col-0, C24 and *fum2*.*2* accessions of Arabidopsis. (a) Maximum capacity for CO_2_ assimilation (*P*_*max*_) was measured at 20 °C and under light- and CO_2_-saturating conditions on adult plants grown and at 20 °C and then exposed to 5, 10, 15, 20 25, or 30 °C treatments for one week. (b-c) In-cabinet measurements of CO_2_ assimilation of plants grown in control conditions (white), and after one day (cyan or orange) or 7 days (blue or red) of cold (b) or warm (c) treatment. Mean standard errors of 4 biological replicates are shown. Different letters above the error bars indicate statistically different values (Analysis of variance, Turkey’s test with a confidence level of 0.95).

**Figure 3:**
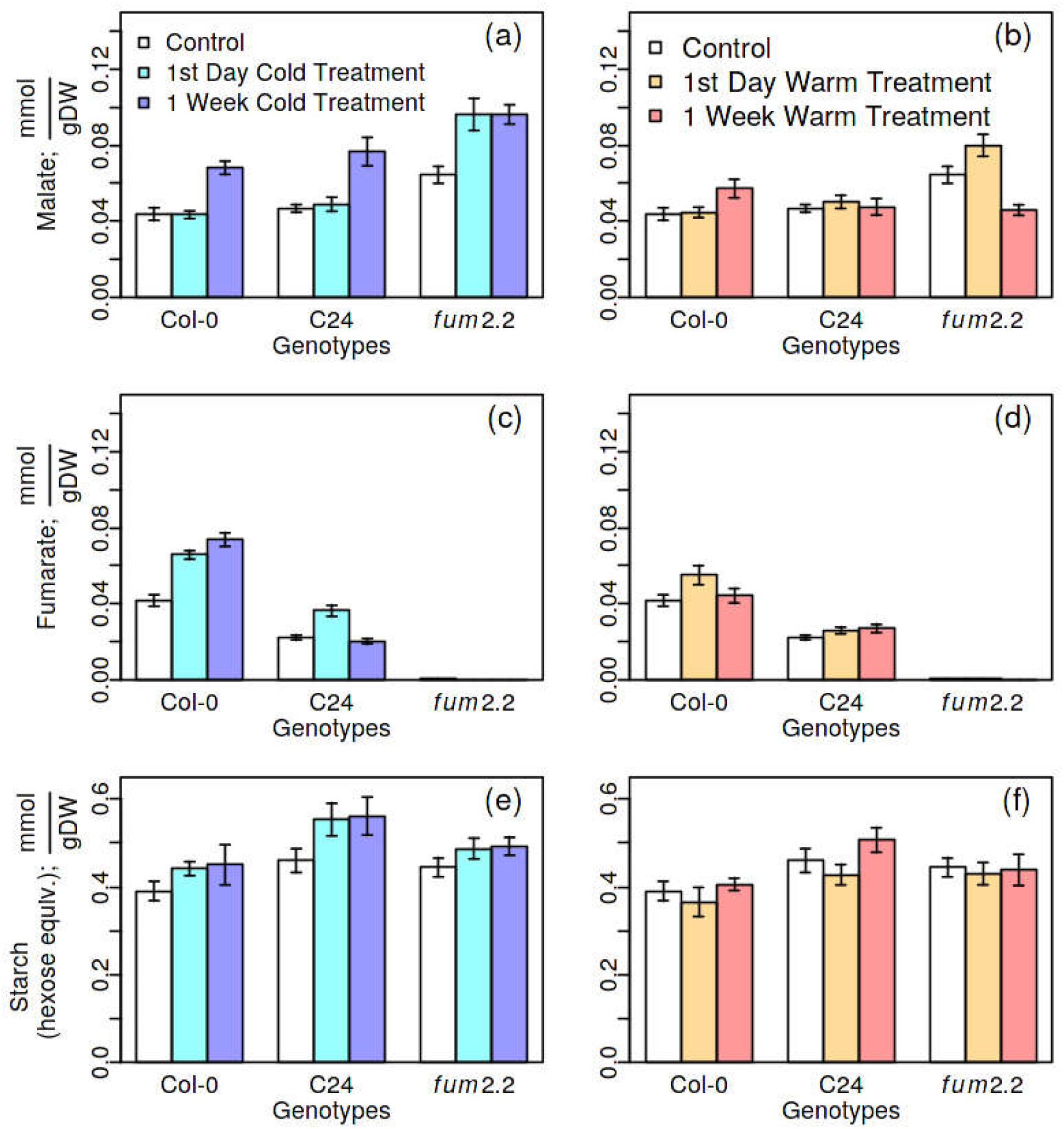
Diurnal accumulation of malate (a-b), fumarate (c-d) and starch shown in hexose equivalents (e-f) in Col-0, C24, and *fum2*.*2* plants under control conditions (white) or after one day (cyan or orange) and seven days (blue or red) of cold (a,c,e) warm (b,d,f) treatments, respectively. Diurnal accumulation was calculated by subtracting the beginning of day concentrations from the end of day concentrations. The uncertainties of 3-4 end of day and beginning of day measurements are shown as mean error bars.

### Data availability

All experimental data, model parameterizations and network analyses are available on Github (https://github.com/HAHerrmann/FMEA_KineticModel) and under the following Zenodo DOI: 10.5281/zenodo.3596623.

## Results

### Temperature effects on photosynthetic acclimation and carbon accumulation

Three Arabidopsis genotypes, Col-0, C24, and *fum2* were grown at a control temperature of 20 °C for 8 weeks. Plants were then either kept at 20 °C or transferred to 5, 10, 15, 25 or 30 °C for 7 days. Maximum photosynthetic capacity (*P*_*max*_) was measured at 20 °C and under light- and CO_2_-saturating conditions on the 1^st^ (Fig. S4) and 7^th^ (Fig. 2a) day of treatment. *P*_*max*_ provides an indication of the acclimation state of the photosynthetic apparatus. *P*_*max*_ did not change on the 1^st^ day of temperature treatments nor did it vary significantly between genotypes (Fig. S4). However, after one week of treatment, Col-0 showed a significant change in *P*_*max*_ in response to the 5, 10, and 30 °C treatments (Fig. 2a). Similarly, the C24 accession showed a change in *P*_*max*_ at 5 and 30 *°C*. The *fum2* mutant, which lacks cytosolic fumarase, does not show a consistent trend in altering its *P*_*max*_ in response to the temperature treatments. The 5 °C treatment is hereafter referred to as cold treatment whilst the 30 °C treatment is referred to as a warm treatment.

CO_2_ fixation under growth conditions was evaluated on the 1^st^ and 7^th^ day of cold (Fig. 2b) and warm treatments (Fig. 2c). In plants transferred to cold, CO_2_ fixation was initially reduced in all three genotypes. However, in Col-0 and C24 plants, photosynthesis recovered after one week of acclimation, reaching the same rate of rate as seen in plants maintained at 20 °C. In contrast, the *fum2* mutant failed to recover its rate of photosynthesis in response to cold. Under warm treatment, Col-0 plants showed lower rates of photosynthesis after one week, whereas C24 and *fum2* plants maintain a constant rate of photosynthesis for the duration of the treatment (Fig. 3c). Rates of respiration show a similar trend to those of photosynthesis in response to cold (Fig. S1a). In the warm treatment, only the *fum2* mutant shows a sustained decrease in respiration after one week (Fig. S1b).

### Partitioning of assimilated carbon between alternative sinks changes in response to both high and low temperature

Measurements of the three major carbon sinks in Arabidopsis leaves - starch, malate, and fumarate - were taken in control conditions, and on the 1^st^ and the 7^th^ day of cold and warm treatment, respectively (Fig. 3). To measure the carbon accumulated during the photoperiod, the beginning of day concentrations were subtracted from the end of day concentrations for each carbon sink. Each of these accumulates in an approximately linear fashion across the 8-hour photoperiod under controlled environment conditions (Dyson et al., 2016). On the 1^st^ day of treatment, diurnal accumulation of malate was unaffected by either cold or warm in Col-0 and C24 plants (Fig. 3a,b; Fig. S5a-c). Fumarate accumulation, however, increased in response to both treatments in these genotypes (Fig. 3c,d; Fig. S5d-f), this response being most pronounced in the cold and in the Col-0 genotype. Consistent with previous published data (Riewe *et al*., 2016), C24 accumulated less fumarate than Col-0 in all conditions, however it accumulated similar amounts of malate. The *fum2* mutant did not accumulate fumarate but increased its malate accumulation on the 1^st^ day of cold and warm treatment. Organic acid accumulation continued to increase in the cold over the week but not in the warm (Fig. 3; Fig. S6). All three genotypes increased their diurnal starch production in response to cold on the 1^st^ day and maintained that increase on the 7^th^ day (Fig. 3e). In the warm, starch accumulation did not change significantly, except in C24 plants on the 7^th^ day of treatment, where it increased (Fig. 3f).

### Reliability engineering identifies cytosolic fumarase as a fail-safe in the metabolic system

The above results and Figures S5 and S6 show that there is not a simple linear relationship between diurnal leaf carbon accumulation and temperature. Using an existing genome-scale metabolic model of Arabidopsis carbon metabolism in the leaf, we have taken a network topology approach to understand how organic acid accumulation changes with temperature treatment. By adapting a Failure Mode and Effect Analysis (FMEA), as outlined in the Materials and Methods, we were able to identify metabolites that pose a high risk to the metabolic system.

High-risk metabolites are those which, in the event of failure, will create the greatest disturbance to the metabolic system. In engineering, the risk factor is calculated based on the probability of failure (*P*_*M*_) and the severity of failure (*S*_*M*_). Here, we estimated *P*_*M*_ based on the length of the shortest path and the number of paths that lead to a metabolite *M. S*_*M*_ can be associated with the number of neighbours of a metabolite and the number of paths that lead to a metabolite (see Materials and Methods). The latter is calculated as the betweenness centrality (i.e. the number of shortest paths that pass through that metabolite).

Table 1 shows the metabolites with the 10 highest risk factors. The average risk factor of all metabolites was 0.005. Malate, oxaloacetate and glucose-6-phosphate stand out as high-risk metabolites in the cytosol (Table 1). In the chloroplast, α-ketoglutarate, glyceraldehyde 3-phosphate, pyruvate, fructose 6-phosphate, ribose 5-phosphate, glucose 6-phosphate, and phosphoenolpyruvate have a high factor. Metabolites related to carbon metabolism and found in the mitochondria have an average or below average risk factor. Only the chloroplast, mitochrondrion, cytosol and peroxisomes compartments were taken into consideration, as specified in the Arnold and Nikoloski (2014) model. The full list of risk factor calculations are available on Zenodo and Github (DOI: 10.5281/zenodo.3596623.

**Table 1:** The ten most high-risk metabolites as identified by the Failure Mode and Effect Analysis (FMEA). An existing metabolic model (Arnold and Nikoloski, 2014; Herrmann *et al*., 2019c) was converted to a graph of carbon metabolism as outlined in the methods. Metabolites with the highest risk factors (*R*) and their cellular locations are shown. Possible cellular locations, as written in the genome-scale metabolic model, include the chloroplast, mitochondrion, cytosol and peroxisomes. *R=S*_*M*_ × *P*_*M*_, where *S*_*M*_*=C*_*M*_ × *N*_*M*_, was calculated such that the severity is equal to the normalized number of neighbours (*N*_*M*_) times the normalized betweenness centrality (*C*_*M*_). PM is based on the length of the shortest path and the number of shortest paths that lead to a metabolite *M*.

| Metabolite | Location | $N_M$ | $C_M$ | $P_M$ | $R \times 10^{-2}$ |
| --- | --- | --- | --- | --- | --- |
| Malate | Cytosol | 1.00 | 0.0374 | 0.500 | 1.87 |
| Oxaloacetate | Cytosol | 1.00 | 0.0564 | 0.429 | 1.42 |
| α-Ketoglutarate | Chloroplast | 0.944 | 0.124 | 0.357 | 4.19 |
| Glyceraldehyde 3-phosphate | Chloroplast | 0.833 | 0.134 | 0.286 | 3.24 |
| Pyruvate | Chloroplast | 0.778 | 0.206 | 0.357 | 5.71 |
| Fructose 6-phosphate | Chloroplast | 0.667 | 0.0685 | 0.357 | 1.63 |
| Glucose 6-phosphate | Cytosol | 0.500 | 0.0695 | 0.500 | 1.74 |
| Ribose 5-phosphate | Chloroplast | 0.500 | 0.0976 | 0.357 | 1.74 |
| Glucose 6-phosphate | Chloroplast | 0.389 | 0.0727 | 0.429 | 1.21 |
| Phosphoenolpyruvate | Chloroplast | 0.389 | 0.129 | 0.286 | 5.71 |

The network structure that connects all high-risk metabolites shows that these are well-connected nodes and are located only a few reactions upstream of a carbon sink (Table 1, Fig. 4). Carbon sinks are generally low-risk metabolites (*R* < 0.001). In particular, fumarate is an endpoint in the system and has a null betweenness centrality (*C*_*M*_). Therefore, changing the concentration of fumarate has a negligible effect on the overall metabolic system. Because changes in the concentration of high-risk metabolites could have a negative effect on system functionality, we hypothesized that adjusting the flux to the nearest carbon sink could minimize the overall system disturbance. Thus, for instance, fumarate could act as a fail-safe, buffering changes in malate, oxaloacetate and other upstream metabolites under different temperature conditions (Fig. 4). To test this hypothesis, we adopted a kinetic modelling approach.

**Figure 4:**
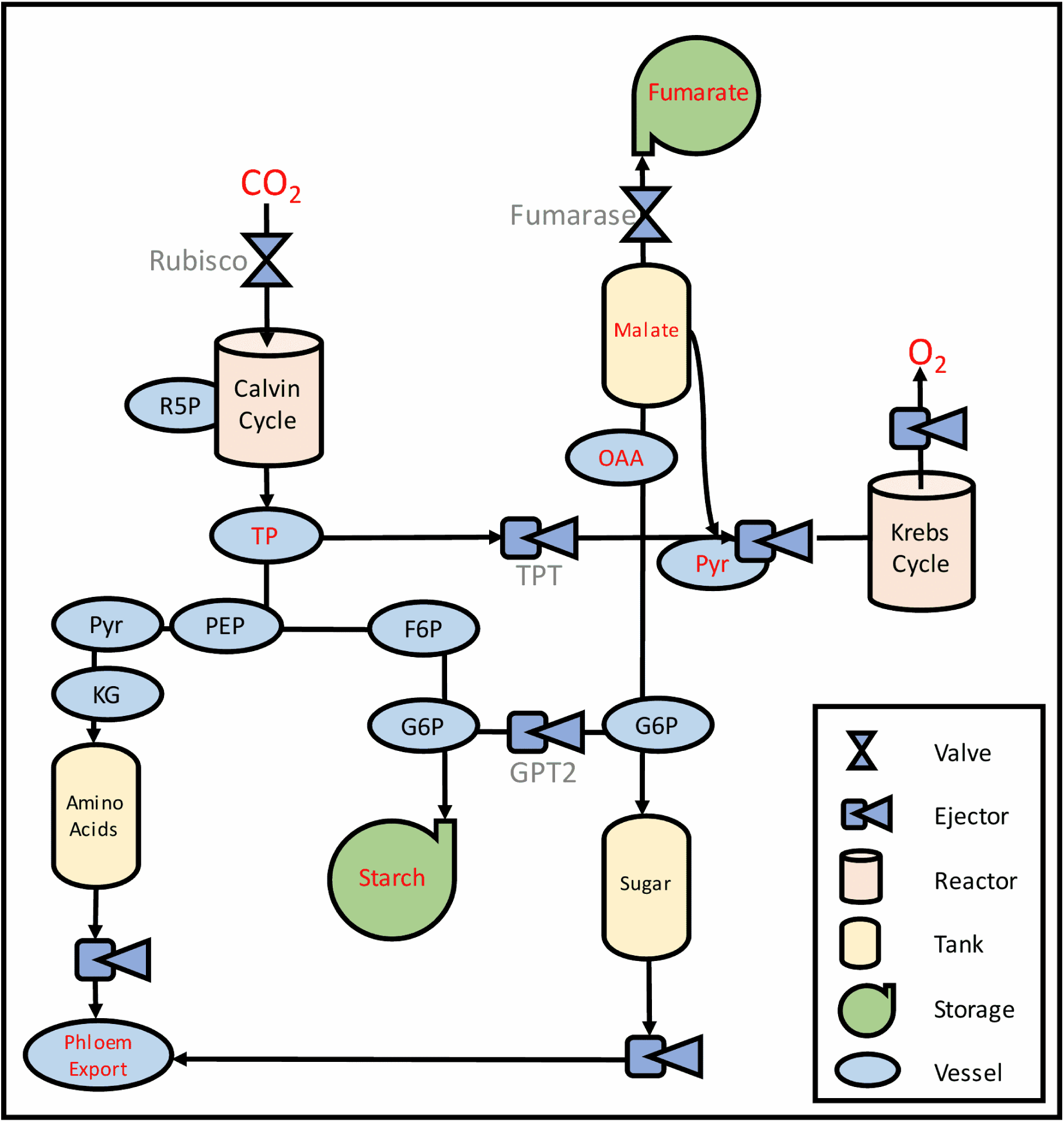
A simplified engineering system of plant primary metabolism connecting high-risk metabolites and their carbon sinks. All high-risk metabolites, as identified in Table 1 were included: R5P (ribose-5-phosphate), TP (glyceraldehyde 3-phosphate), PEP (phosphoenolpyruvate), Pyr (pyruvate), KG (α-ketoglutarate), F6P (fructose 6-phosphate), G6P (glucose 6-phosphate), OAA (Oxaloacetate) and Malate. Only branch and end point metabolites were included in the kinetic model and are shown in red.

### Organic acid accumulation across temperatures requires no regulatory mechanisms

To further test whether fumarate acts as a buffer for malate across the physiological temperature range of Arabidopsis, we constructed a kinetic model of the primary reactions in leaf carbon metabolism (Fig. 1, Table S1). To keep the number of reactions in this model to a minimum, only relevant branch and end points (as shown in red in Fig. 4) were included in the model. In comparison to starch, malate and fumarate, neither sugars nor amino acids accumulate substantially in Arabidopsis leaves in our growth conditions (Dyson *et al*., 2016); however, it must be assumed that there is a significant flux through these pools that constitutes phloem export (marked as C export in Fig. 1; Lalonde *et al*., 2003, Ainsworth *et al*., 2011). Rates of photosynthesis and respiration (Fig. 2b,c; Fig. S1) were used to constrain the model. The remaining parameters were estimated to fit the measured malate, fumarate and starch concentrations in control conditions, and on the first day of cold and warm treatment, respectively. It is assumed that on the first day of temperature treatment, plants will not yet have significantly acclimated to the stress condition; thus, observed changes in metabolism would be mainly the result of kinetic or regulatory effects rather than changes in enzyme concentrations.

When fitting separate models for Col-0, *fum2* and C24 plants, the models predict the same kinetic parameters for all reactions except for the conversion of fumarate to malate and that of triose phosphate to starch (Table 2). C24 has significantly lower cytosolic fumarase activity than Col-0 and accumulates only about three quarters of the amount of fumarate accumulated in Col-0 leaves (Riewe *et al*., 2016). The models predict no cytosolic fumarase activity in *fum2* and a reduced activity in C24 compared to Col-0 plants, confirming experimental evidence. The results obtained show that increased fumarate production in response to both warm and cold treatment can result of temperature-dependent enzyme kinetics rather than post-translational regulation of enzyme activity. While we cannot exclude that these regulatory effects are occurring, the model simply shows that a solution for organic acid production without temperature-dependent regulatory effects is possible.

**Table 2:**
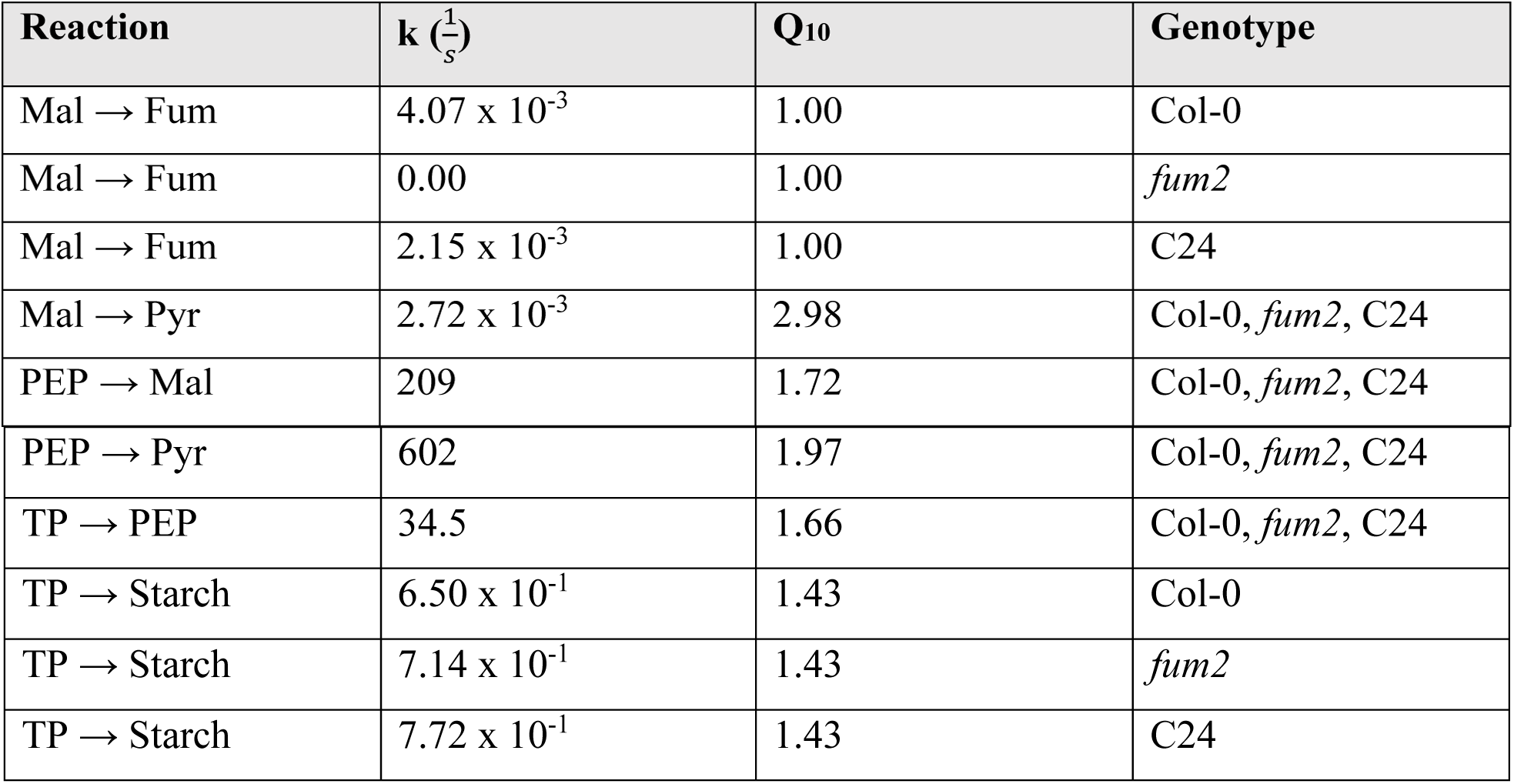
Parameters for a kinetic model of carbon metabolism. The metabolites malate (Mal), fumarate (Fum), pyruvate (Pyr), phosphoenolpyruvate (PEP), triose phosphate (TP) were used to construct the model. The full table including the rates of photosynthesis and respiration set as model constraints are shown in the Supplementary Materials (Table **S1**).

| Reaction | $k \left(\frac{1}{s}\right)$ | $Q_{10}$ | Genotype |
| --- | --- | --- | --- |
| Mal → Fum | $4.07 \times 10^{-3}$ | 1.00 | Col-0 |
| Mal → Fum | 0.00 | 1.00 | *fum2* |
| Mal → Fum | $2.15 \times 10^{-3}$ | 1.00 | C24 |
| Mal → Pyr | $2.72 \times 10^{-3}$ | 2.98 | Col-0, *fum2*, C24 |
| PEP → Mal | 209 | 1.72 | Col-0, *fum2*, C24 |
| PEP → Pyr | 602 | 1.97 | Col-0, *fum2*, C24 |
| TP → PEP | 34.5 | 1.66 | Col-0, *fum2*, C24 |
| TP → Starch | $6.50 \times 10^{-1}$ | 1.43 | Col-0 |
| TP → Starch | $7.14 \times 10^{-1}$ | 1.43 | *fum2* |
| TP → Starch | $7.72 \times 10^{-1}$ | 1.43 | C24 |

In the solutions to the models, most effective *Q*_*10*_ values are estimated to be around 2.0 (Table 2), as is typical for temperature-dependence of biological reactions (Elias, 2014). The cytosolic fumarase reaction, however, shows no apparent temperature-dependence, effective *Q*_*10*_ = 1.0. Although we are aware that all enzymatic reactions are affected by temperature, we note that an effective *Q*_*10*_ = 1.0 does not mean that the reaction is not temperature-sensitive but rather that the corresponding enzyme is not limiting. Thus, our model results merely demonstrate that differences in the rate constants are enough to explain the observed difference between genotypes. The conversion of malate to pyruvate is, according to the models, the most temperature-sensitive reaction; however, it carries little overall flux. The conversions of phosphoenolpyruvate to malate and pyruvate, on the other hand, carry a substantial flux and show a strong temperature-dependence. The counter-play of these two reactions, along with the observed changes in rates of photosynthesis and respiration, are sufficient to account for the experimentally observed changes in organic acid concentrations under warm and cold temperature treatments.

The rate of export was allowed to adjust freely when fitting the model. This rate accounts for the total remaining carbon, which is assumed to be exported from the leaves, primarily in the form of sugars, but will include other forms of fixed carbon retained in the leaf (Wilkinson and Douglas, 2003). All genotypes are predicted to decrease their diurnal rates of export on the first day of cold treatment (Fig. 5). On the first day of warm treatment, Col-0 and *fum2* plants show little change in the rate of export, compared to control conditions (Fig. 5). C24 plants, however, are predicted to increase their rates of export under warm treatment. This is the result of decreased starch accumulation (Fig. 3f) as well as decreased rates of respiration (Fig. S1) in response to warm treatment.

**Figure 5:**
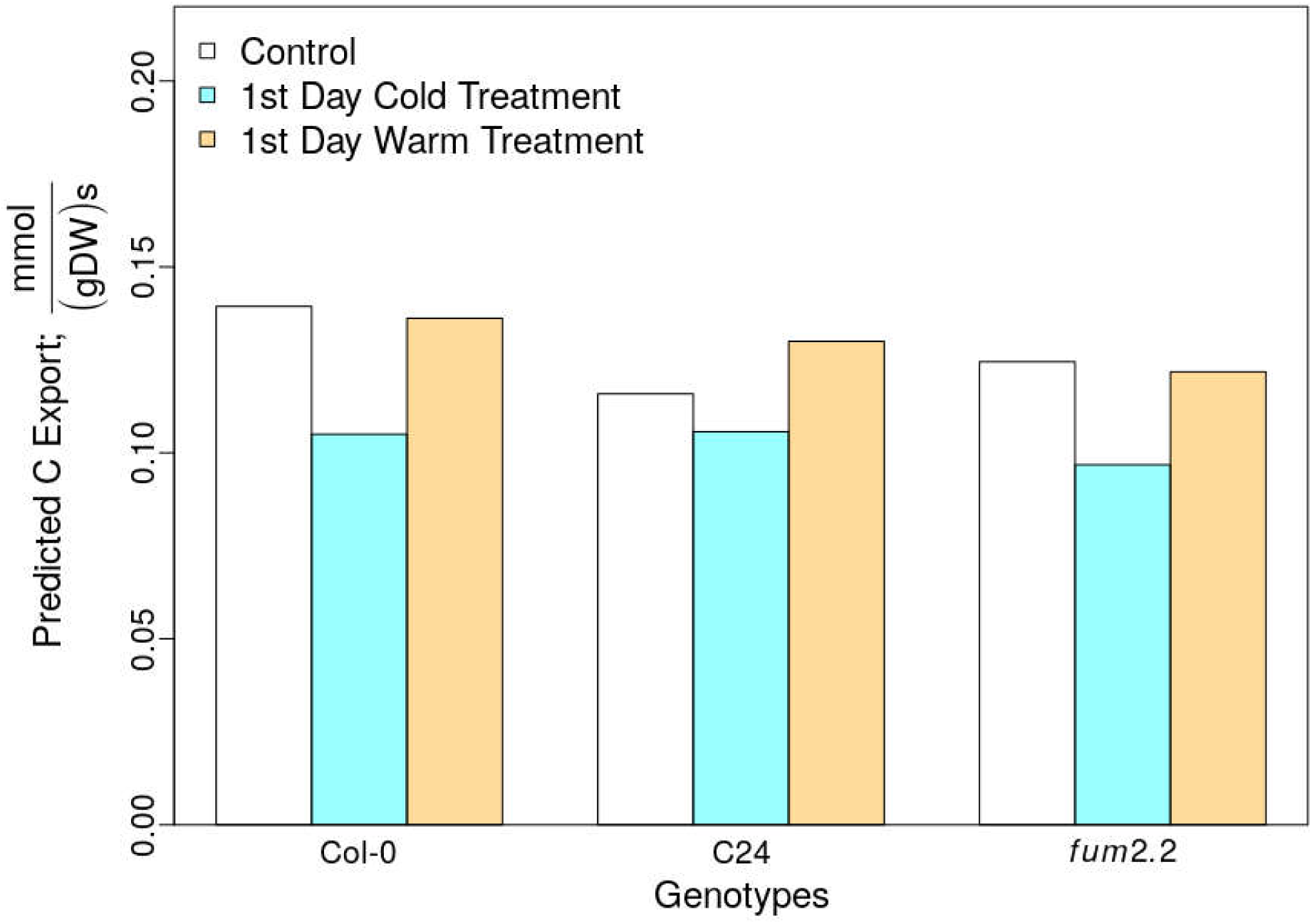
Diurnal export rates as predicted from kinetic modelling. Rates of photosynthesis and respiration were used to constrain the model and the remaining model parameters were estimated to fit the measured metabolite concentrations (Table S1, Fig’s. 1, 2, S1). Export rates estimated for control conditions (white), the first day of cold (cyan) and the first day of warm (orange) treatment, represent the remaining carbon not stored in the leaf.

## Discussion

Understanding plant responses to short-term changes in the environment is of considerable importance if we wish to improve crop yields in the light of changing climates. The relationship between these environmental changes and the signal transduction pathways which promote cellular adjustment is still poorly understood. Because all of metabolism is affected by temperature, it is difficult to pinpoint an obvious temperature-sensor or to assess the direct effect that metabolic changes have on gene expression (Herrmann *et al*., 2019a). For this reason, rather than looking at individual components of a system, it is important to look at the entire system and consider the role that individual components play within it (Kitano, 2002). In this sense, formal frameworks, which conceptualize, rather than merely describe, the functioning of its components, are required (Lazebnik, 2003).

Robustness can be used to assess how well a system maintains function across changing environmental conditions. Robustness, however, does not capture the likelihood that the system will be able to perform its function independently of the environmental conditions. This is captured by measures of reliability.

Here, we have adapted the failure mode and effect analysis (FMEA) framework used in reliability engineering and applied it to plant carbon metabolism. We used a metabolic graph of carbon metabolism to identify metabolites which can be considered of high risk to the system, due to their high probability of failure and severity of impact on the rest of the system. From an engineering perspective, high-risk components should be carefully controlled and regulated such that they can maintain their functionality in the event of failure. A fail-safe, for example, can be used to ensure that perturbations are mitigated effectively. Such a fail-safe increases the overall reliability of a system by increasing its likelihood of correct functioning. A good fail-safe is able to mitigate many different types of system failure, and fail-safes are of particular importance under abrupt and unexpected changes.

Diurnal fumarate accumulation increases in response to both cold and warm treatment (Fig’s 3, S5). This surprising observation is consistent with flux to fumarate acting as an inherent fail-safe to the metabolic system. Our FMEA analysis identifies malate and oxaloacetate, the immediate precursors of fumarate, as high-risk components. Fumarate itself is of low risk to the metabolic system and changing its concentration has negligible effect on overall system functionality. An influx of carbon can be re-directed to fumarate without disturbing the system. Our results show that, unlike fumarate, the amount of malate accumulating through the day in both wild type accessions is surprisingly constant in response to initial warm and cold treatment and only changes following acclimation. Mathematical modelling confirms that an increased accumulation of fumarate and a constant accumulation of malate are metabolically plausible under the Arrhenius law (Arrhenius, 1889), and that no regulatory mechanisms are required for fumarate accumulation to increase at both high and low temperatures. Nevertheless, active regulation of fumarate accumulation under these conditions cannot be excluded.

To date, there is little direct experimental evidence for the presence of a cytosolic fumarase enzyme in plant species other than *Arabidopsis thaliana* (Chia et al. 2000). However, a recent phylogenetic study demonstrated that several close relatives of *A. thaliana* possess orthologues of the *fum2* gene (Zubimendi *et al*., 2018). This study also shows that other plant species obtained the gene through parallel evolution. Because fumarate has recently evolved in Arabidopsis species and relatives (Zubimendi *et al*., 2018), it is likely that other mechanisms regulate the concentration of high-risk metabolites (like malate) in other species. For instance, phosphoenolpyruvate carboxylase (PEPC), which produces malate from oxaloacetate, is inhibited by malate across many plant species (O’Leary *et al*., 2011). This negative feedback loop could have the same effect as a fumarate sink, in that it maintains a constant accumulation of malate, even when the carbon influx is changing. This regulatory mechanism, however, is intricately dependent on phosphorylation and cellular pH (O’Leary *et al*., 2011), both of which may also be affected by changing environmental conditions. Thus, the evolution of a cytosolic fumarase fail-safe provides an alternative control mechanism regulating the malate concentration, while at the same time maintaining metabolic fluxes and allowing efficient storage of fixed carbon.

Photosynthetic acclimation to cold is dependent on the presence of FUM2 activity or protein (Dyson *et al*., 2016). Here, we have shown that C24, which has reduced FUM2 content has an attenuated the acclimation response. Furthermore, we show that FUM2 is also essential for acclimation of photosynthetic capacity to high temperatures. Photosynthetic capacity, measured under control temperature and CO_2_ and light-saturating conditions, provides a readily measurable indicator of the acclimation state of the photosynthetic apparatus (Herrmann *et al*., 2019b). Mutant *fum2* plants are unable to accumulate fumarate and do not show a consistent adjustment of *P*_*max*_ in response to temperature. C24 plants accumulate intermediate levels of fumarate and attenuated temperature acclimation of *P*_*max*_, compared with Col-0. The *P*_*max*_ acclimation responses of Col-0 and C24 highlight that, although the changes in CO_2_ assimilation in growth conditions may be modest, there is a significant metabolic response associated with these changes.

Although malate accumulation is buffered on the first day of temperature treatment, it does increase in response to sustained temperature treatment, as part of the acclimation response. Our FMEA analysis suggests that malate concentration should be tightly regulated to support a specific system functionality. This functionality may change in response to cold conditions, at which point a different rate of malate accumulation may be required. Either way, the required concentration is likely to be controlled by adjusting the flux to the low-risk metabolite fumarate.

The functionality of a metabolic system may vary under changing environmental conditions and the ability to achieve function should be maximised. Thus, finding reliable rather than robust traits, should be of equal if not of higher priority in optimizing a metabolic system. Methods for capturing system reliability currently remain underutilized in biology.

While we have focused here on malate as a high-risk metabolite, our analysis has identified other high-risk candidates. Interestingly, some of them have previously been noted as critical components of acclimation (Timm *et al*., 2012; Dyson *et al*., 2015; Weise *et al*., 2019). For example, it was shown that expression of GTP2, a chloroplast glucose 6-phosphate/phosphate translocator, is required for acclimation to high light in the Arabidopsis accession Wassilewskija-4 (Dyson *et al*., 2015). Our FMEA highlights glucose 6-phosphate in the chloroplast and in the cytosol as high-risk components, which are upstream of the starch and sucrose carbon sinks in the metabolic network. GPT2 allows for appropriate distribution of carbon between these two sinks. When this control mechanism is broken, the two glucose 6-phosphate metabolite concentrations cannot be regulated and acclimation to high light is affected (Dyson *et al*., 2015). Other high-risk metabolites identified by our FMEA, such as glyceraldehyde 3-phosphate, pyruvate, phosphoenolpyruvate, and α-ketoglutarate, may also play important roles in maintaining system reliability under changing environmental conditions.

In conclusion, we were able to show that cytosolic fumarase, an enzyme promoting the accumulation of fumarate, a metabolite with few known metabolic functions, can act as a fail-safe, preventing over-accumulation of the high-risk metabolite malate. We have used FMEA to discuss alterations in plant carbon metabolism under different temperature conditions, and we propose FMEA as a tool to assess the reliability of biological systems.

Our proposed FMEA framework is computationally inexpensive and can be applied to all of metabolism in order to identify pathways that are especially important for maintaining system reliability. As we have shown FMEA allows for the quantification of high-risk and low-risk components in existing systems. However, FMEA could also be used to quantify the risk associated with metabolic alterations that are the result of gene deletions or insertions. For this we would recommend a gene-centric rather than a reaction-centric view of the network that underlies the metabolic system of interest. This approach holds the potential to be used in metabolic engineering, for the study of synthetic pathways and the impact of pathways alterations on system reliability.

## Acknowledgements

HAH is supported by a Biotechnology and Biological Sciences Research Council (BBSRC) Doctoral Training Partnership stipend (BB/M011208/1) and PIC is supported by BBSRC research grant (BB/S009078/1). We thank Armida-Irene Gjindali for proof-reading the final manuscript.

## Author Contributions

HAH, JMS and GNJ conceived the study. HAH conducted the experiments and developed the FMEA framework. PIC contributed to the analysis of the data. All authors co-wrote the manuscript.

## Supplementary Materials

**Figure S1:**
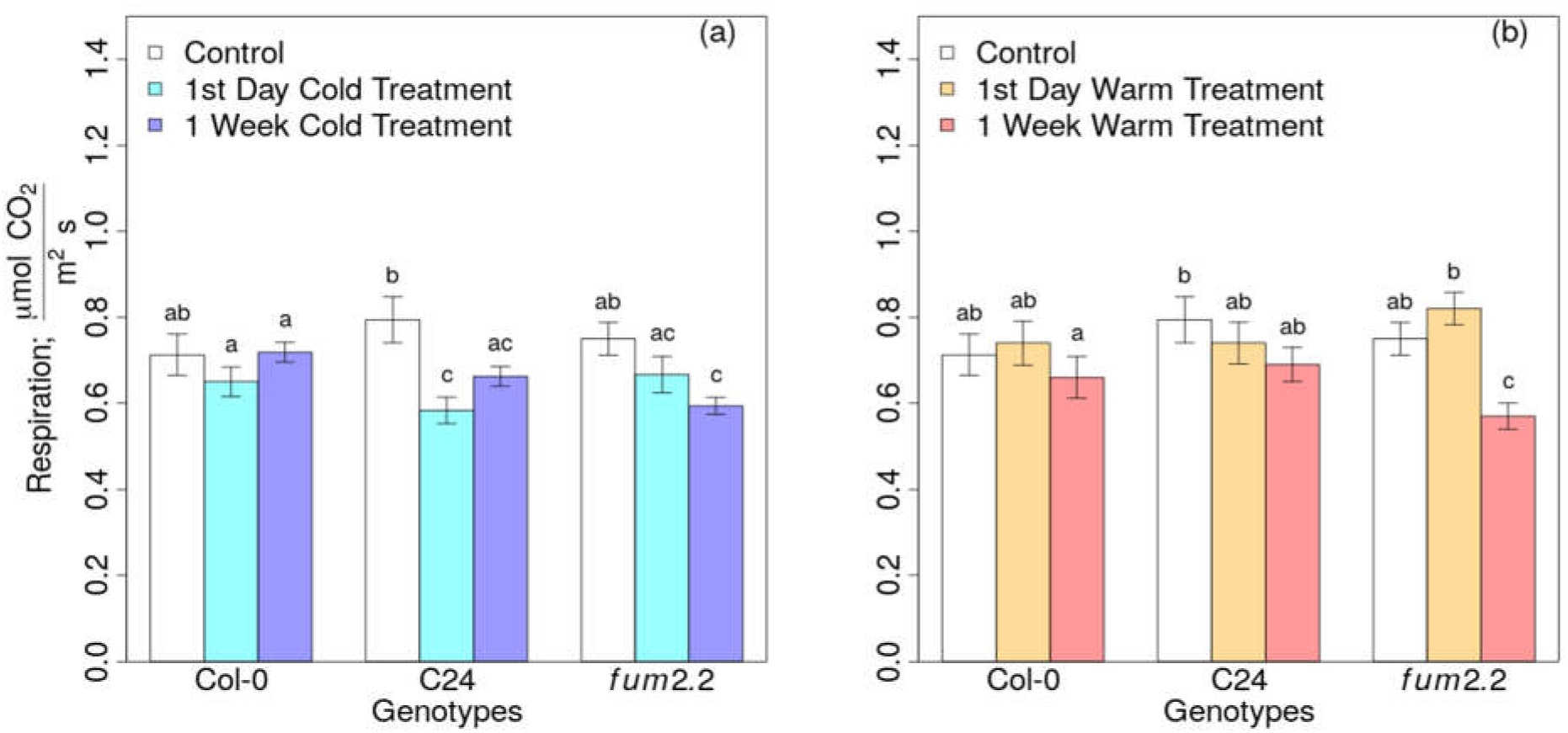
Respiration measurements during the first and seventh day of the cold (a) and warm treatments (b). CO_2_ respiration (R) measurements were taken in the dark in adult Arabidopsis plants grown under control temperature conditions (white), after one or seven days of cold treatment (cyan and blue) and warm treatment (orange and red), respectively. Standard mean errors of 3-4 biological replicates are shown. Different letters above the error bars indicate statistically different values (Analysis of variance, Turkey’s test with a confidence level of 0.95).

**Figure S2:**
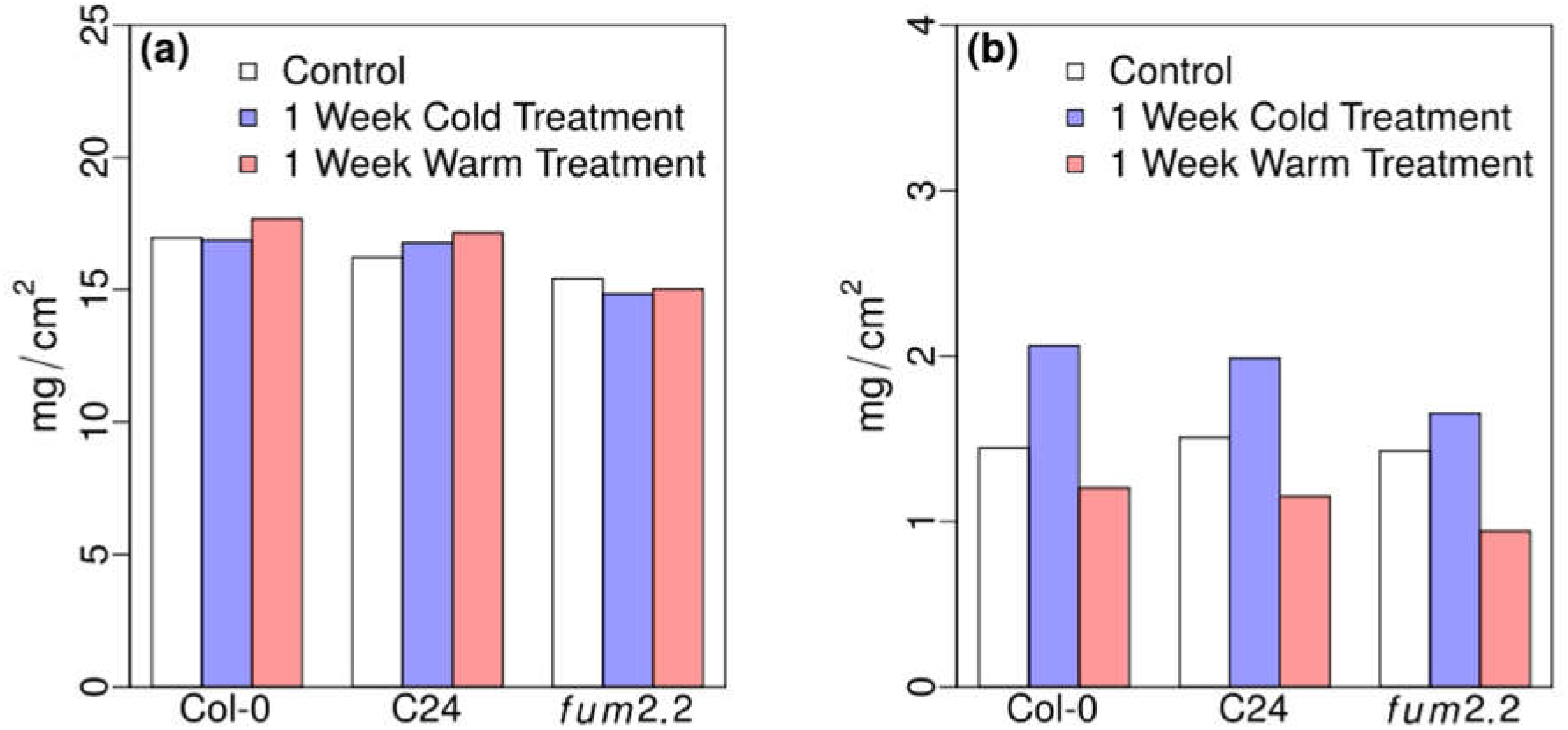
Leaf area to weight comparisons. Arabidopsis leaf fresh weight (a) and dry weight (b) per unit area measured in adult plants under control conditions (20 *°C*), cold stress (5 *°C*) or warm stress (30 *°C*) treatment. Col-0 values are shown in gray, *fum2* are shown in black and C24 in white. Each measurement is based on 4 biological replicates. Ratios of the averages are shown.

**Figure S3:**
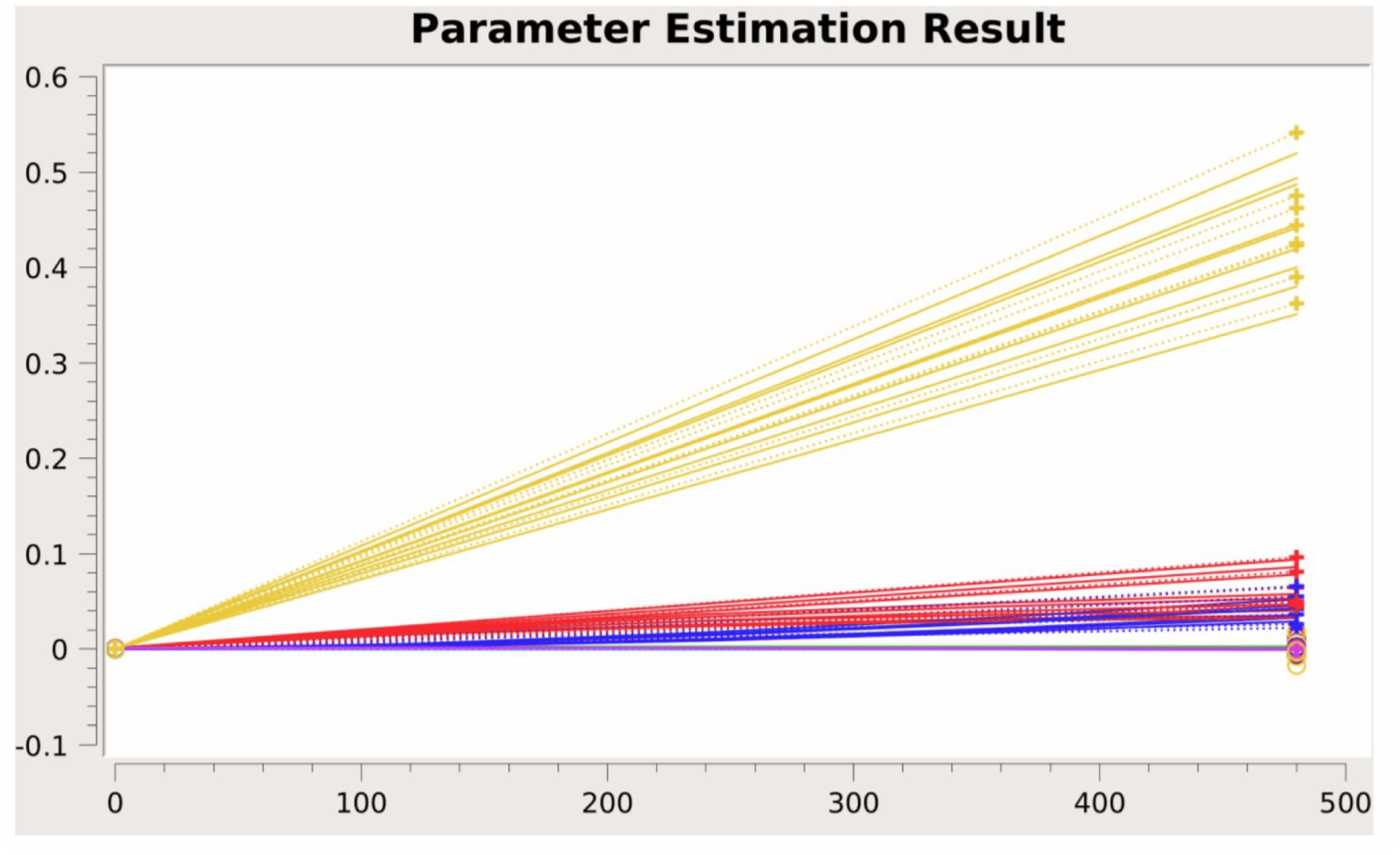
Screenshot of the parameter estimation of metabolites concentrations (*M/(gFW)*, y-axis) over time (*s*, x-axis) as done in COPASI (Version 4.27.217). Parameters were fitted to the experimentally measured beginning and end of day metabolite concentrations of the kinetic model using Hooke & Jeeves algorithm with an iteration limit of 500, a tolerance of 10^−8^ and a Rho of 0.2. Fits were done according to measurements taken under control conditions, on the first day of cold and warm treatment for starch (yellow), malate (red), and fumarate (blue). All other metabolites concentrations were kept between 5.0e-5 and 1.4e-4 *M/(gFW*). The experimental data is shown as crosses joined by dotted lines; the fitted data is shown as straight lines.

**Figure S4:**
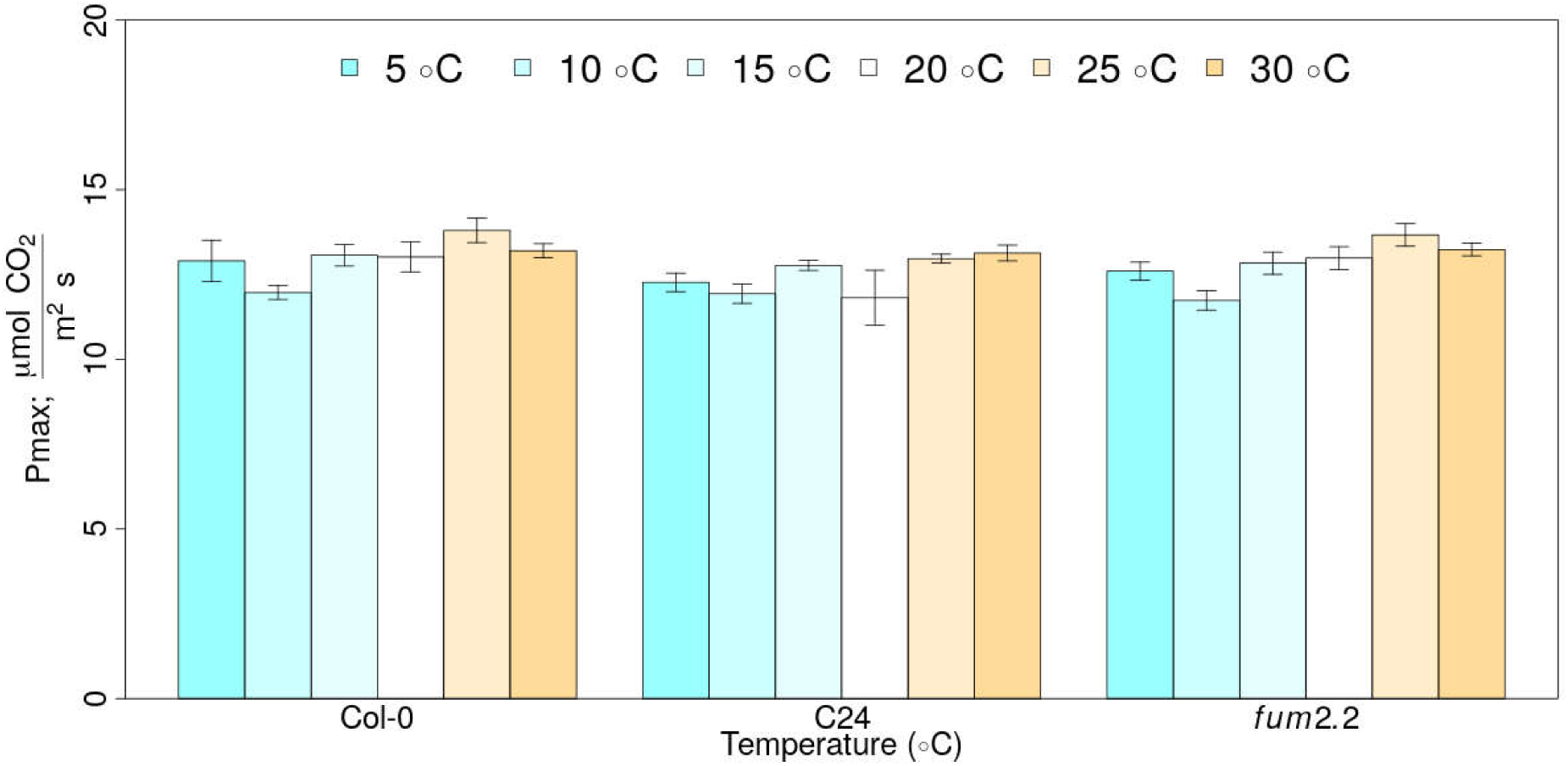
Measurements of the maximum photosynthetic capacity (*P*_*max*_) on the first day of temperature treatments. CO_2_ assimilation was measured under light- and CO_2_-saturating conditions in control plants kept at 20 *°C* and in plants subjected to 6 hours of 5, 10, 15, 25, or 30 *°C* temperature treatment. Standard mean errors of 4 biological replicates are shown.

**Figure S5:**
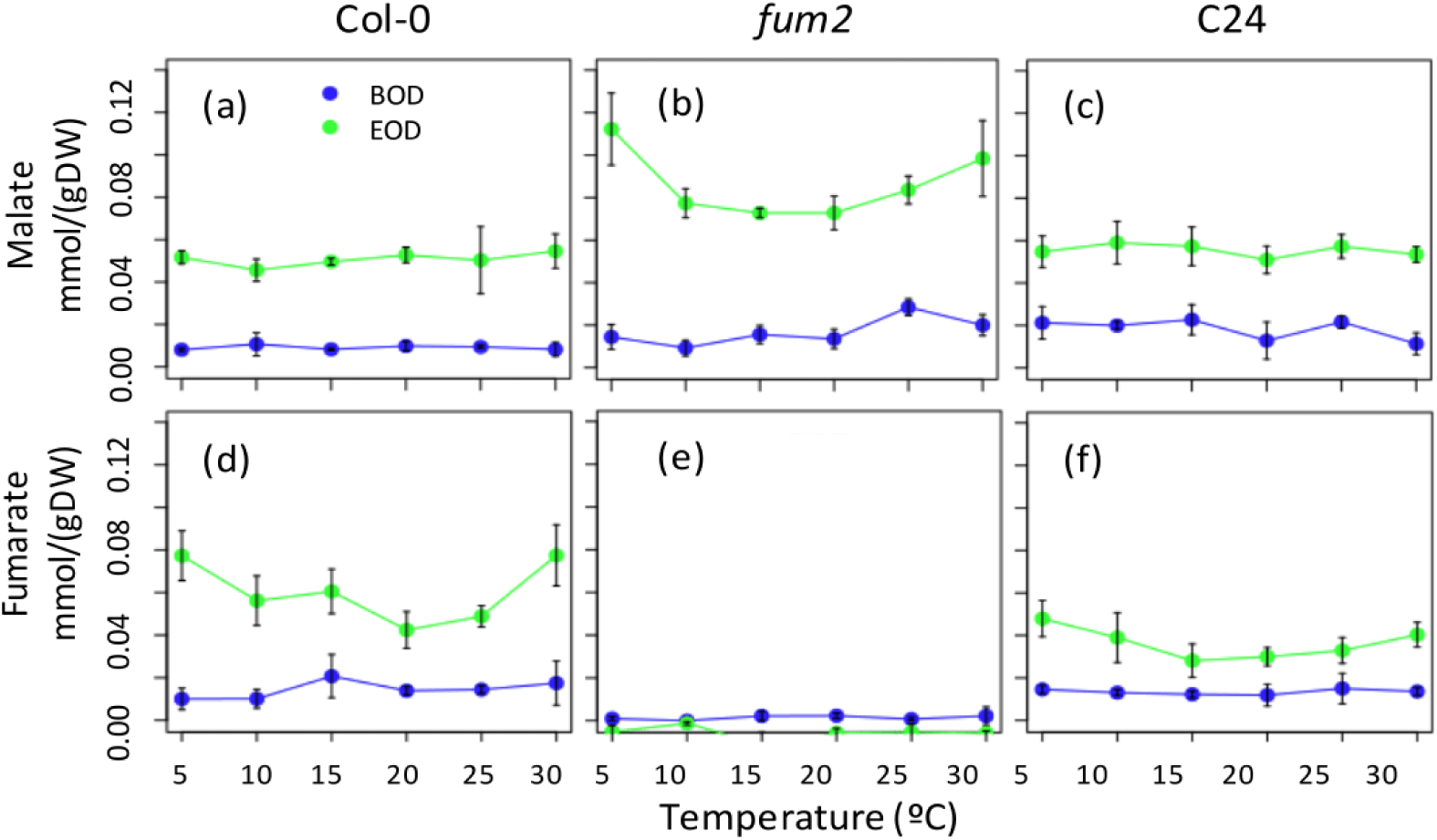
Organic acid concentrations of three Arabidopsis genotypes measured across 6 temperatures on the first day of treatment. Beginning of day (BOD; blue) and end of day (EOD; green) concentration of malate and fumarate are shown for Col-0 (a,d), *fum2* (b,e) and C24 (c,f) genotypes. Standard mean errors of 3-4 biological replicates are shown.

**Figure S6:**
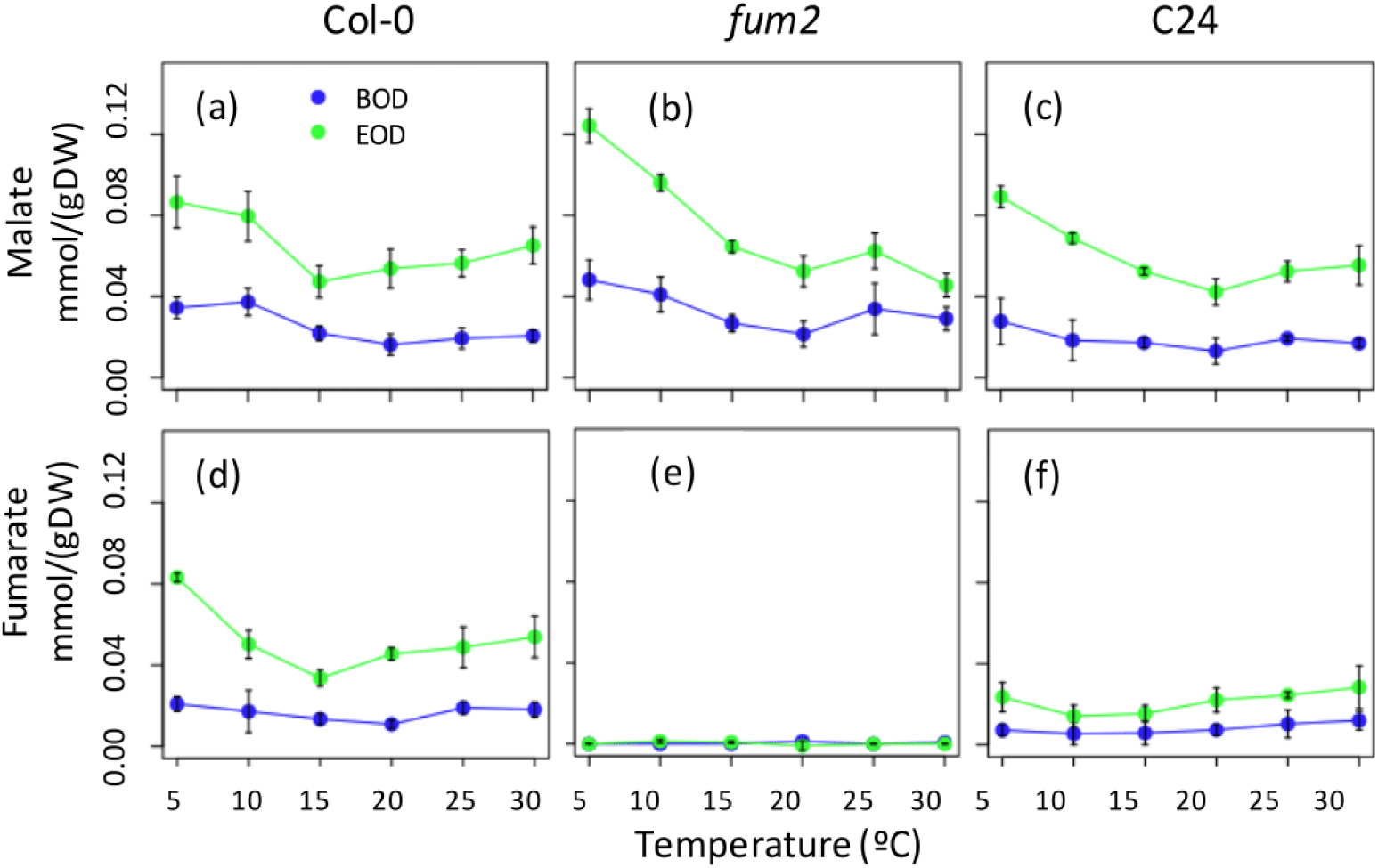
Organic acid concentrations of three Arabidopsis genotypes measured across 6 temperatures on the seventh day of treatment. Beginning of day (BOD; blue) and end of day (EOD; green) concentration of malate and fumarate are shown Col-0 (a,d), *fum2* (b,e) and C24 (c,f) genotypes. Standard mean errors of 3-4 biological replicates are shown.

**Table S1:**
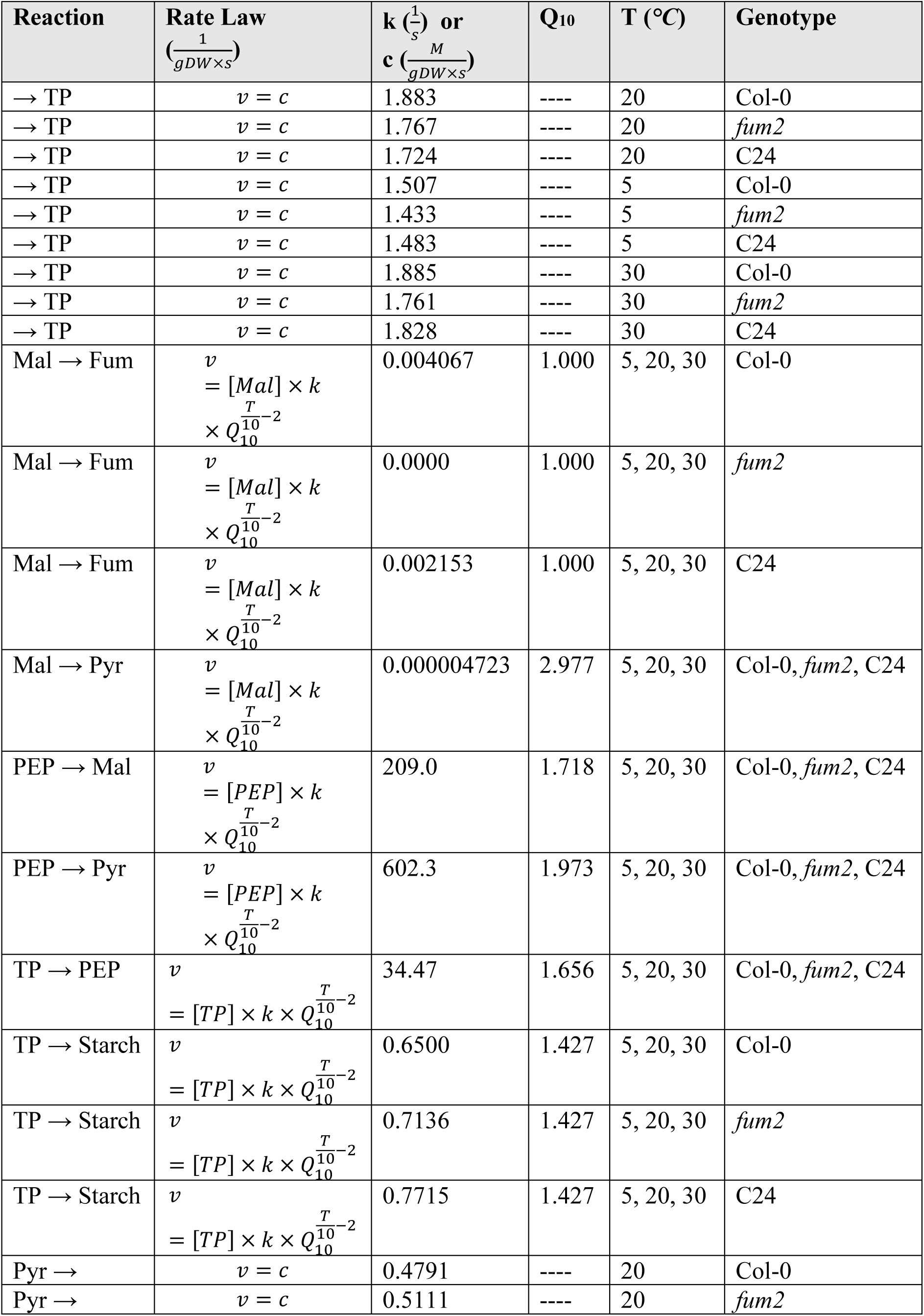
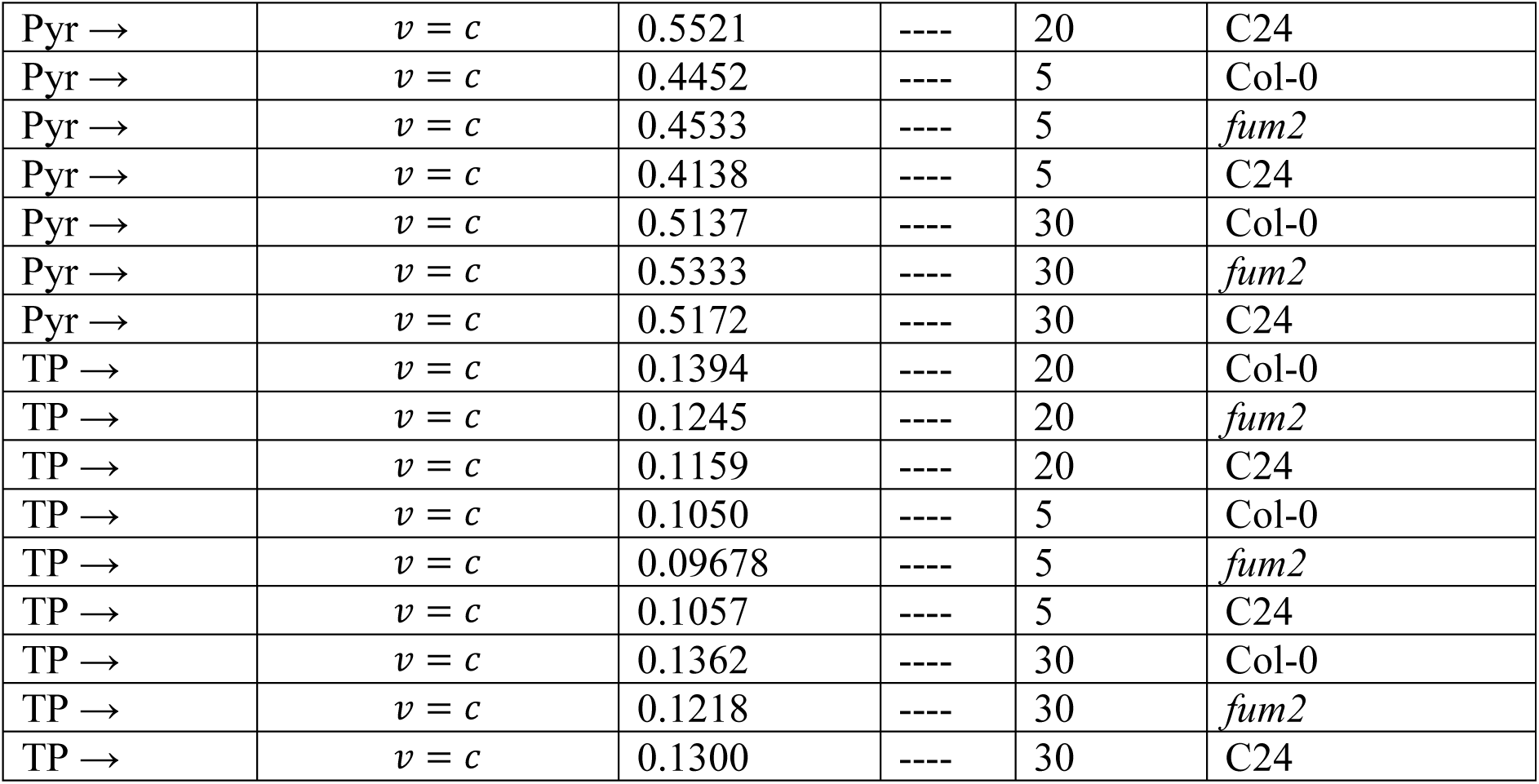
Reaction rate laws for a small kinetic model of carbon metabolism. Reaction rates used to set up and parametrize a kinetic model. The same abbreviations as in Fig. **3** apply, in addition to malate (Mal) and fumarate (Fum).

## References

Abegg R. (1899). Untersuchungen über die Chemischen Affinitäten. Abhandlungen aus den Jahren 1864, 1867, 1879. Leipzig: Wilhelm Engelmann.

Ainsworth E.A. & Bush D.R. (2011) Carbohydrate export from the leaf: A highly regulated process and target to enhance photosynthesis and productivity. Plant Physiol. 155, 64–69.

Athanasiou K., Dyson B.C., Webster R.E & Johnson G.N. (2010). Dynamic acclimation of photosynthesis increases plant fitness in changing environments. Plant Physiol. 152, 366–373.

Arnold A. & Nikoloski Z. (2014). Bottom-up reconstruction of Arabidopsis and its application to determining the metabolic costs of enzyme production. Plant Physiol. 165, 1380–1391.

Arrhenius S.A. (1889). Über die Dissociationswärme und den Einfluß der Temperatur auf den Dissociationsgrad der Elektrolyte. Physikal. Chemie 4, 96–116.

Bakera J.W., Schubert M. & Faberb M.H. (2008). On the assessment of robustness. Structural Safety 30, 253–267.

Beck J.B., Schmuths H. & Schaal B.A. (2008). Native range genetic variation in Arabidopsis thaliana is strongly geographically structured and reflects Pleistocene glacial dynamics. Mol. Ecol. 17, 902–915.

Changnon S.A., Pielke R.A.Jr., Changnon D., Sylves R.T. & Pulwarty R. (2000). Human factors explain the increased losses from weather and climate extremes. Americ. Meterol. Soc. 81, 437–442.

Chia D.W., Yoder T.J., Reiter W.-D. & Gibson S.I. (2000). Fumaric acid: an overlooked form of fixed carbon in Arabidopsis. Planta 211, 743–751.

Christensen B. & Nielsen J. (2000). Metabolic network analysis. A powerful tool in metabolic engineering. Adv. Biochem. Eng. Biotechnol. 66, 209–231.

Ding Y., Shi Y. & Yang S. (2020). Molecular regulation of plant responses to environmental temperatures. Mol. Plant. 13, 544–564.

Dyson B.C., Allwood J.W., Feil R., Xu Y., Miller M., Bowsher C.G., Goodacre R., Lunn J.E. & Johnson G.N. (2015). Acclimation of metabolism to light in Arabidopsis thaliana: the glucose 6-phosphate/phosphate translocator GPT2 directs metabolic acclimation. Plant, Cell & Environ. 38, 1404–1417.

Dyson B.C., Miller M.A.E., Feil R., Rattray N., Bowsher C.G., Goodacre R., Lunn J.E. & Johnson G.N. (2016). FUM2, a cytosolic fumarase, is essential for acclimation to low temperature in Arabidopsis thaliana. Plant Physiol. 172, 118–127.

Elias M., Wieczorek G., Rosenne S. & Tawfik D.S. (2014). The universality of enzymatic rate-temperature dependency. Trends Biochem. Sci. 39, 1–7.

Faust K., Dupont P., Callut J., van Helden J. (2010). Pathway discovery in metabolic networks by subgraph extraction. Bioinformatics 26, 1211–1218.

Fernie A.R. & Martinoia, E. (2009). Malate Jack of all trades or master of a few? Phytochem. 70, 828–832.

Frainay C. & Jourdan F. (2017). Computational methods to identify metabolic sub-networks based on metabolomic profiles. Brief. Bioinform. 18, 43–56.

Freeman L.C. (1977). A Set of Measures of Centrality Based on Betweenness. Sociometry 40, 35–41.

Gerstl M.P., Klamt S., Jungreuthmayer C. & Zanghellini J. (2016). Exact quantification of cellular robustness in genome-scale metabolic networks. Bioinformatics. 32, 730–737.

Gnan S., Priest A., Kover P.X. (2014). The genetic basis of natural variation in seed size and seed number and their trade-off using Arabidopsis thaliana MAGIC lines. Genetics 198, 1751–1758.

Herrmann H.A., Schwartz J.-M. & Johnson G.N. (2019a). Metabolic acclimation—a key to enhancing photosynthesis in changing environments? J. of Exp. Bot. 12: 3043–3056.

Herrmann H.A., Schwartz J.-M. & Johnson, G.N. (2019b). From empirical to theoretical models of light response curves-linking photosynthetic and metabolic acclimation. Photosynth. Res. doi: 10.1007/s11120-019-00681-2.

Herrmann H.A., Dyson B.C., Vass L., Johnson G.N. & Schwartz J.-M. (2019c). Flux sampling is a powerful tool to study metabolism under changing environmental conditions. npj Sys. Biol. & App. 3, 32.

Hikosaka K., Ishikawa K., Borhigidai A., Muller O. & Onoda Y. (2006). Temperature acclimation of photosynthesis: mechanisms involved in the changes in temperature dependence of photosynthetic rate. J. of Exp. Bot. 57, 291–302.

Holme P. (2011). Metabolic Robustness and Network Modularity: A Model Study. PLoS One 6, e16605.

Hooke R. & Jeeves T. (1961) Direct search solutions of numerical and statistical problems. J. of the Assoc. for Computing Machinery. 8, 212–229.

Hurry V., Strand Å., Furbank R. & Stitt M. (2000). The role of inorganic phosphate in the development of freezing tolerance and the acclimatization of photosynthesis to low temperature is revealed by the pho mutants of Arabidopsis thaliana. The Plant J. 24, 383–396.

Jeong H., Tombor B., Albert R., Oltvai Z.N. & Barabasi A.L. (2000). The large-scale organization of metabolic networks. Nature 407, 651–654.

Jeong H., Mason S.P., Barabasi A.L. & Oltvai Z.N. (2001). Lethality and centrality in protein networks. Nature 411, 41–42.

Johnson, G.N. & Murchie E. (2011). Gas exchange measurements for determination of photosynthetic efficiency in Arabidopsis leaves. Method. in Molec. Biol. 775, 311–326.

Kaplan F., Kopka J., Haskell D.W., Zhao W., Schiller K.C., Gatzke N., Sung D.Y. & Guy C.L. (2004). Exploring the temperature-stress metabolome of Arabidopsis. Plant Physiol. 136, 4159–4168.

King J.P. & Jewett W.S. (2010). Robustness Development and Reliability Growth: Value Adding Strategies for New Products and Processes. Pearson Education, Technology & Engineering.

Kitano H. (2002). Systems biology: A brief overview. Science 295, 1662–1664.

Kitano H. (2004). Biological Robustness. Nat. Rev. Genetics 5, 826–837.

Kovermann P., Meyer S., Hortensteiner S., Picco C., Scholz-Starke J., Ravera S., Lee Y., Martinoia E. (2007) The Arabidopsis vacuolar malate channel is a member of the ALMT family. Plant J. 52, 1169–1180.

Lalonde S., Tegeder M., Throne-Holst M., Frommer W.B. & Patrick J.W. (2003) Phloem loading and unloading of sugars and amino acids. Plant, Cell & Environm. 26, 37–56.

Lazebnik Y. (2003). Can a biologist fix a radio? – Or, what I learned while studying apoptosis. Cancer Cell 2, 179–182.

Lesk C., Rowhani P. & Ramankutty N. (2016). Framing the way to relate climate extremes to climate change. Nature 529, 84–87.

Lewis N.E., Nagarajan H. & Palsson B.O. (2012). Constraining the metabolic genotype-phenotype relationship using a phylogeny of in silico methods. Nat. Rev. Microbiol., 10, 291–305.

O’Leary B., Park J. & Plaxton W.C. (2011). The remarkable diversity of plant PEPC (phosphoenolpyruvate carboxylase): recent insights into the physiological functions and post-translational controls of non-photosynthetic PEPCs. Biochem J. 436, 15–34.

Powell J.P. & Reinhard S. (1993). Photosynthesis, photoinhibition and low temperature acclimation in cold tolerant plants. Photosyn. Res. 37, 19–39.

Powell J.P. & Reinhard S. (2016). Measuring the effects of extreme weather events on yields. Weather Clim. Extr. 12, 69–79.

Pracharoenwattana I., Zhou W., Keech O., Francisco P.B., Udomchalothorn T., Tschoep H., Stitt M., Gibon Y. & Smith S.M. (2010). Arabidopsis has a cytosolic fumarase required for the massive allocation of photosynthate into fumaric acid and for rapid plant growth on high nitrogen. Plant J. 1, 785–795.

Rausanda M. & Øienb K. (1996). The basic concepts of failure analysis. Reliab. Eng. & Sys. Safety 53, 73–83.

Riewe D., Jeon H.-J., Lisec J., Heuermann M.C., Schmeichel J., Seyfarth M., Meyer R.C., Willmitzer L. & Altmann T. (2016). A naturally occurring promoter polymorphism of the Arabidopsis FUM2 gene causes expression variation and is associated with metabolic and growth traits. The Plant J. 88, 826–838.

Saa P.A. & Nielsen L.K. (2017) Formulation, construction and analysis of kinetic models of metabolism: A review of modelling frameworks. Biotechnol. Adv. 35, 981–1003.

Sadras V.O. (2007). Evolutionary aspects of the trade-off between seed size and number in crops. Field Crops Research 100, 125–138.

Scott I.M., Ward J.L., Miller S.J. & Beale M.H. (2014). Opposite variations in fumarate and malate dominate metabolic phenotypes of Arabidopsis salicylate mutants with abnormal biomass under chilling. Physiol. Plant. 88, 660–674.

Smith A. & Stitt M. (2007). Coordination of carbon supply and plant growth. Plant, Cell, & Environ. 30, 1126–1149.

Stamatis D.H. (1995). Failure Mode and Effect Analysis: FMEA from Theory to Execution. ASQC Quality Press 53, Milwaukee, Wisc.

Stitt M. & Hurry V.A. (2002). plant for all seasons: alterations in photosynthetic carbon metabolism during cold acclimation in Arabidopsis. Curr. Opinion Plant Biol. 5, 199–206.

Strand Å., Foyer C.H., Gustafsson P., Gardeström P. & Hurry V. (2003). Altering flux through the sucrose biosynthesis pathway in transgenic Arabidopsis thaliana modifies photosynthetic acclimation at low temperatures and the development of freezing tolerance. Plant, Cell, & Env. 26, 523–535.

Streb S. & Zeemana SC. (2012). Starch Metabolism in Arabidopsis. Arabidop. Book 10, e0160.

Timm S., Mielewczick M., Florian A., Frankenback S., Dreissen A., Hocken N., Fernie A.R., Walter A. & Bauwe H. (2012). High-to-Low CO2 acclimation reveals plasticity of the photorespiratory pathway and indicates regulatory links to cellular metabolism of Arabidopsis. PLoS One 7, e42809.

Trenberth K.E. (2012). Framing the way to relate climate extremes to climate change. Climatic Change 115, 283–290.

Webber A.N., Nie G.-Y. & Long S.P. (1994). Acclimation of photosynthetic proteins to rising atmospheric CO2. Photosynth. Res. 39, 413–425.

Weise S.E., Liu T., Childs K.L., Preiser L.P., Katulski H.M., Perrin-Prozondek C. & Sharkey T.D. (2019). Transcriptional regulation of the glucose-6-phosphate/phosphate translocator 2 is related to carbon exchange across the chloroplast envelope. Front. Plant Sci. 10, 827.

Wilkinson T.L. & Douglas AE. (2003). Phloem amino acids and the host plant range of the polyphagous aphid, Aphis fabae. Entomol. Exp. Appl. 106, 103–113.

Yamori W., Hikosaka K. & Way D.A. (2014). Temperature response of photosynthesis in C3, C4, and CAM plants: temperature acclimation and temperature adaptation. Photosyn. Res. 119, 101–117.

Yassine A.A. (2007). Investigating product development process reliability and robustness using simulation. J. of Eng. Design 18, 545–561.

Zell M.B., Fahnenstick H., Maier A., Saigo M., Voznesenskaya E.V., Edwards G.E., Andreo C., Schleifenbaum F., Zell C., Drincovich M.F. & Maurino V.G. (2010). Analysis of Arabidopsis with highly reduced levels of malate and fumarate sheds light on the role of these organic acids as storage molecules. Plant Physiol. 152, 1251–1562.

Zhang Q., Kong X., Yu Q., Ding Y., Li X. & Yang Y. (2019). Responses of PYR/PYL/RCAR ABA receptors to contrasting stresses, warm and cold in Arabidopsis. Plant Signal Behav. 14, 1670596.

Zubimendi J.P., Martinatto A., Valacco M.P., Moreno S., Andreo C.S., Drincovich M.F. & Tronconi M.A. (2018). The complex allosteric and redox regulation of the fumarate hydratase and malate dehydratase reactions of Arabidopsis thaliana Fumarase 1 and 2 gives clues for understanding the massive accumulation of fumarate. FEBS J. 285, 2205–2224.

