## Supplementary Materials for "A Failure Mode and Effect Analysis of plant metabolism reveals why cytosolic fumarase is required for temperature acclimation in Arabidopsis"

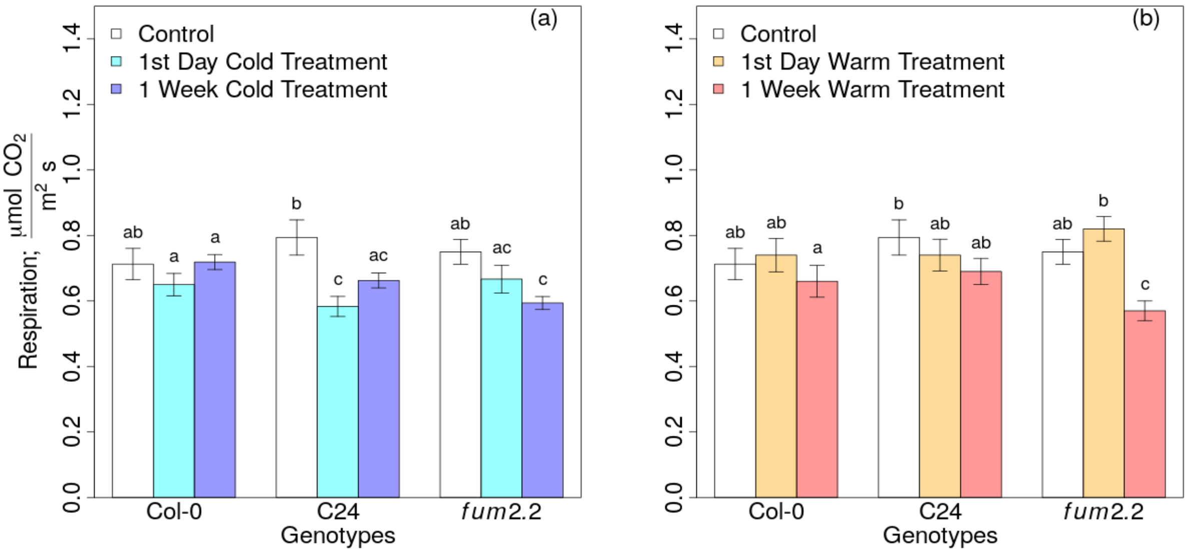


**Figure S1:** Respiration measurements during the first and seventh day of the cold (a) and warm treatments (b). $CO_2$ respiration (R) measurements were taken in the dark in adult Arabidopsis plants grown under control temperature conditions (white), after one or seven days of cold treatment (cyan and blue) and warm treatment (orange and red), respectively. Standard mean errors of 3-4 biological replicates are shown. Different letters above the error bars indicate statistically different values (Analysis of variance, Turkey's test with a confidence level of 0.95).

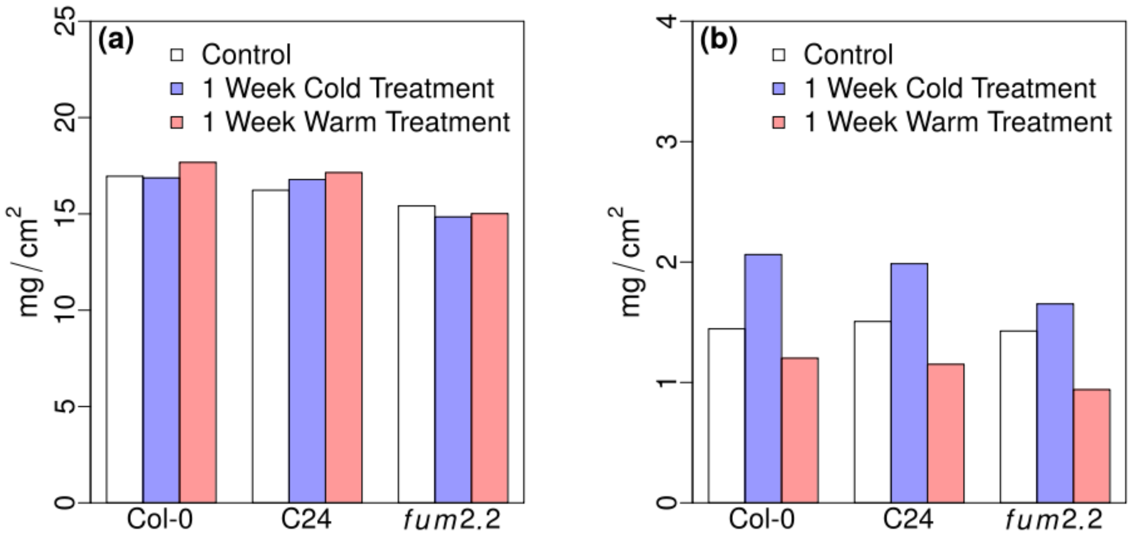


**Figure S2:** Leaf area to weight comparisons. Arabidopsis leaf fresh weight (a) and dry weight (b) per unit area measured in adult plants under control conditions (20 *°C*), cold stress (5 *°C*) or warm stress (30 *°C*) treatment. Col-0 values are shown in gray, *fum2* are shown in black and C24 in white. Each measurement is based on 4 biological replicates. Ratios of the averages are shown.

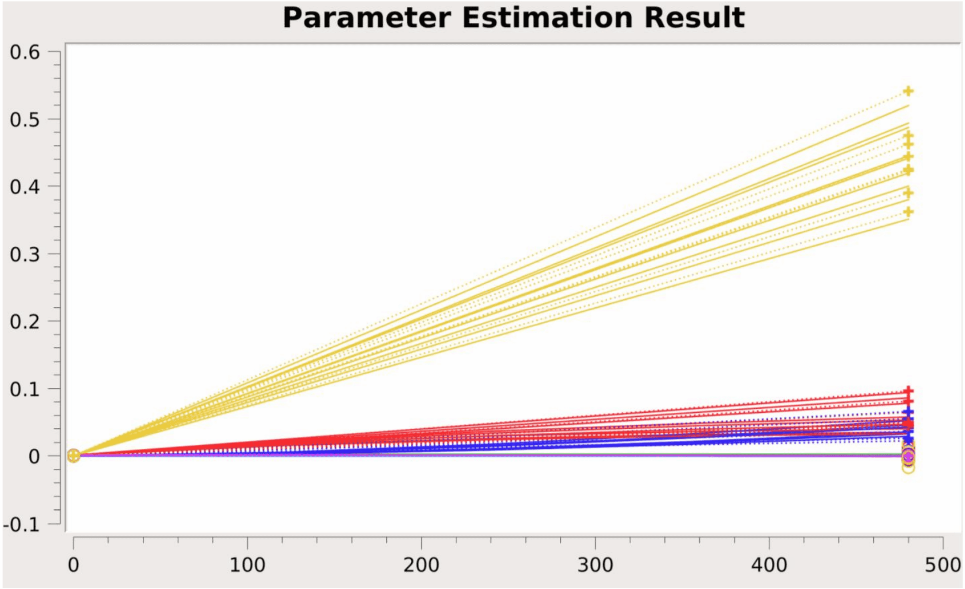


**Figure S3**: Screenshot of the parameter estimation of metabolites concentrations (*M/(gFW)*, y-axis) over time (*s*, x-axis) as done in COPASI (Version 4.27.217). Parameters were fitted to the experimentally measured beginning and end of day metabolite concentrations of the kinetic model using Hooke & Jeeves algorithm with an iteration limit of 500, a tolerance of $10^{-8}$ and a Rho of 0.2. Fits were done according to measurements taken under control conditions, on the first day of cold and warm treatment for starch (yellow), malate (red), and fumarate (blue). All other metabolites concentrations were kept between 5.0e-5 and 1.4e-4 *M/(gFW* ). The experimental data is shown as crosses joined by dotted lines; the fitted data is shown as straight lines.

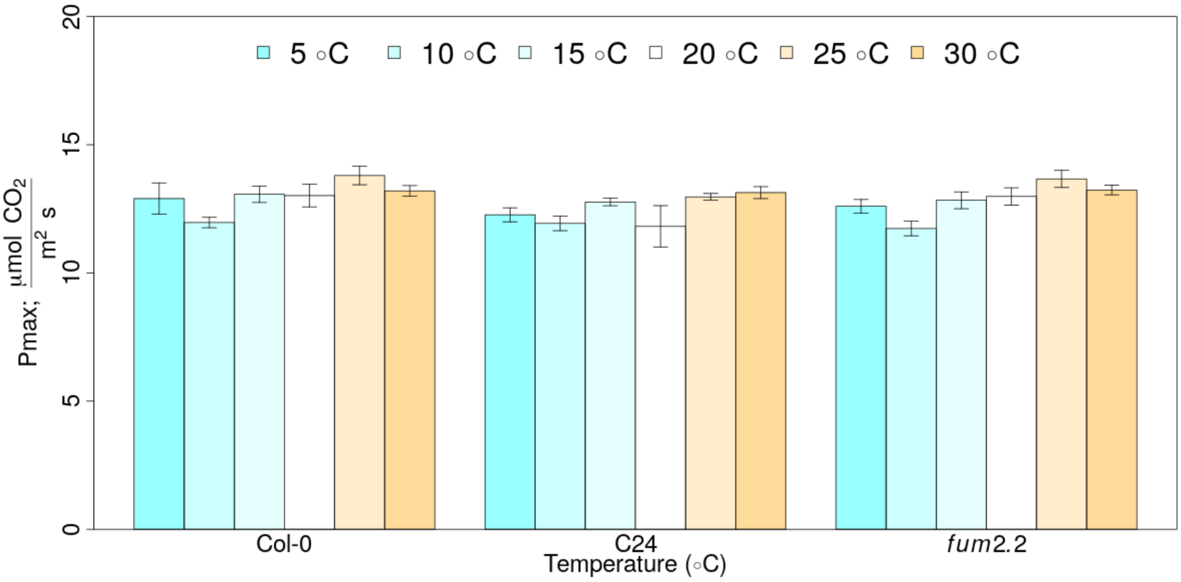


**Figure S4**: Measurements of the maximum photosynthetic capacity ($P_{max}$) on the first day of temperature treatments. $CO_2$ assimilation was measured under light- and $CO_2$-saturating conditions in control plants kept at 20 *°C* and in plants subjected to 6 hours of 5, 10, 15, 25, or 30 *°C* temperature treatment. Standard mean errors of 4 biological replicates are shown.

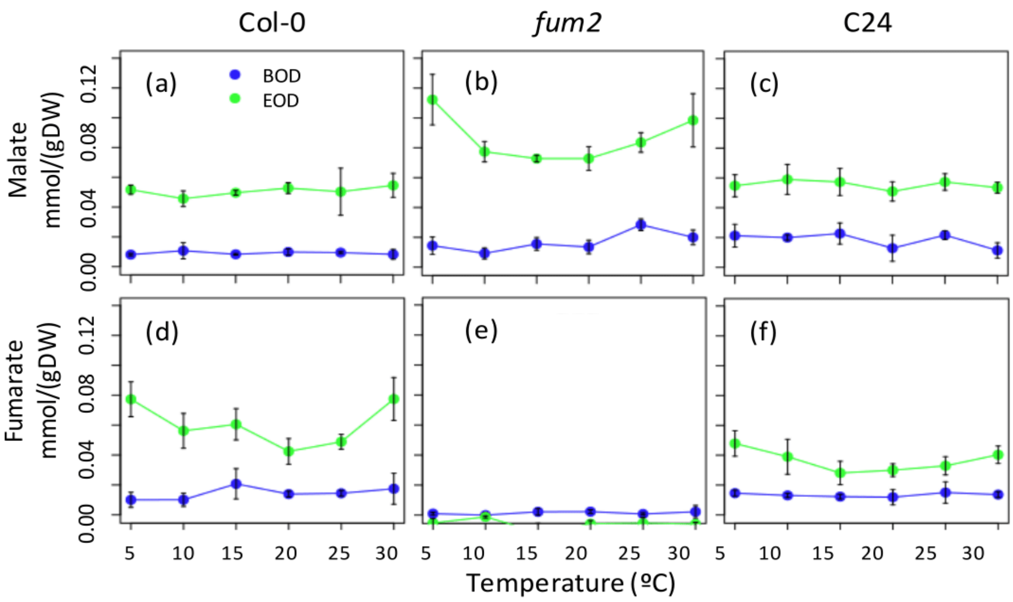


**Figure S5**: Organic acid concentrations of three Arabidopsis genotypes measured across 6 temperatures on the first day of treatment. Beginning of day (BOD; blue) and end of day (EOD; green) concentration of malate and fumarate are shown for Col-0 (a,d), *fum2* (b,e) and C24 (c,f) genotypes. Standard mean errors of 3-4 biological replicates are shown.

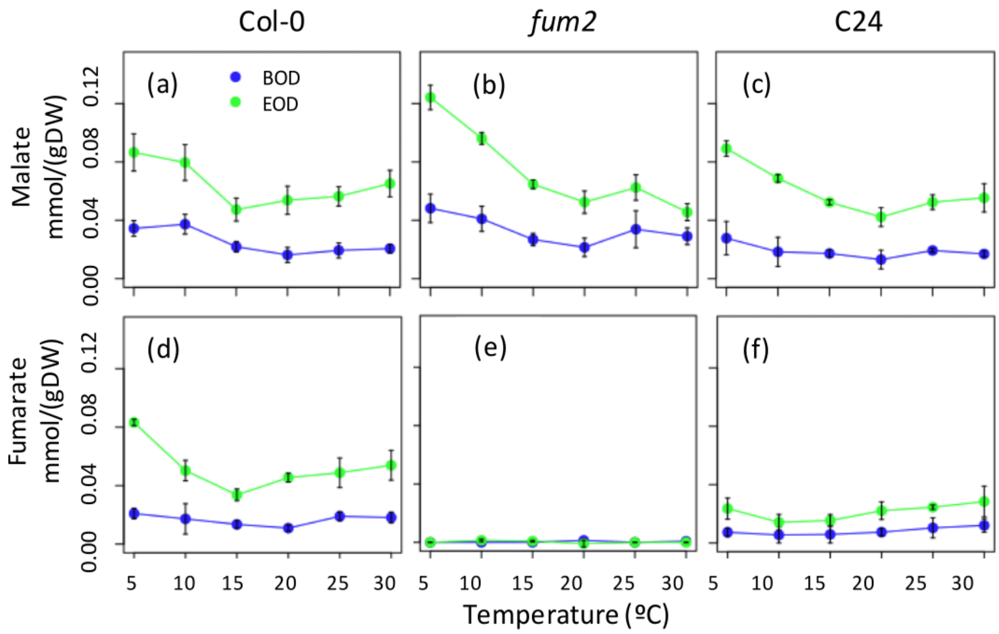


**Figure S6:** Organic acid concentrations of three Arabidopsis genotypes measured across 6 temperatures on the seventh day of treatment. Beginning of day (BOD; blue) and end of day (EOD; green) concentration of malate and fumarate are shown Col-0 (a,d), *fum2* (b,e) and C24 (c,f) genotypes. Standard mean errors of 3-4 biological replicates are shown.

**Table S1:** Reaction rate laws for a small kinetic model of carbon metabolism. Reaction rates used to set up and parametrize a kinetic model. The same abbreviations as in Fig. **3** apply, in addition to malate (Mal) and fumarate (Fum).

| **Reaction** | **Rate Law** $(\frac{1}{g\,DW \times s})$ | **k** $(\frac{1}{s})$ **or c** $(\frac{M}{g\,DW \times s})$ | **Q₁₀** | **T (°C)** | **Genotype** |
|---|---|---|---|---|---|
| → TP | $v = c$ | 1.883 | ---- | 20 | Col-0 |
| → TP | $v = c$ | 1.767 | ---- | 20 | *fum2* |
| → TP | $v = c$ | 1.724 | ---- | 20 | C24 |
| → TP | $v = c$ | 1.507 | ---- | 5 | Col-0 |
| → TP | $v = c$ | 1.433 | ---- | 5 | *fum2* |
| → TP | $v = c$ | 1.483 | ---- | 5 | C24 |
| → TP | $v = c$ | 1.885 | ---- | 30 | Col-0 |
| → TP | $v = c$ | 1.761 | ---- | 30 | *fum2* |
| → TP | $v = c$ | 1.828 | ---- | 30 | C24 |
| Mal → Fum | $v = [Mal] \times k \times Q_{10}^{\frac{T}{10}-2}$ | 0.004067 | 1.000 | 5, 20, 30 | Col-0 |
| Mal → Fum | $v = [Mal] \times k \times Q_{10}^{\frac{T}{10}-2}$ | 0.0000 | 1.000 | 5, 20, 30 | *fum2* |
| Mal → Fum | $v = [Mal] \times k \times Q_{10}^{\frac{T}{10}-2}$ | 0.002153 | 1.000 | 5, 20, 30 | C24 |
| Mal → Pyr | $v = [Mal] \times k \times Q_{10}^{\frac{T}{10}-2}$ | 0.000004723 | 2.977 | 5, 20, 30 | Col-0, *fum2*, C24 |
| PEP → Mal | $v = [PEP] \times k \times Q_{10}^{\frac{T}{10}-2}$ | 209.0 | 1.718 | 5, 20, 30 | Col-0, *fum2*, C24 |
| PEP → Pyr | $v = [PEP] \times k \times Q_{10}^{\frac{T}{10}-2}$ | 602.3 | 1.973 | 5, 20, 30 | Col-0, *fum2*, C24 |
| TP → PEP | $v = [TP] \times k \times Q_{10}^{\frac{T}{10}-2}$ | 34.47 | 1.656 | 5, 20, 30 | Col-0, *fum2*, C24 |
| TP → Starch | $v = [TP] \times k \times Q_{10}^{\frac{T}{10}-2}$ | 0.6500 | 1.427 | 5, 20, 30 | Col-0 |
| TP → Starch | $v = [TP] \times k \times Q_{10}^{\frac{T}{10}-2}$ | 0.7136 | 1.427 | 5, 20, 30 | *fum2* |
| TP → Starch | $v = [TP] \times k \times Q_{10}^{\frac{T}{10}-2}$ | 0.7715 | 1.427 | 5, 20, 30 | C24 |
| Pyr → | $v = c$ | 0.4791 | ---- | 20 | Col-0 |
| Pyr → | $v = c$ | 0.5111 | ---- | 20 | *fum2* |
| Pyr → | $v = c$ | 0.5521 | ---- | 20 | C24 |
| Pyr → | $v = c$ | 0.4452 | ---- | 5 | Col-0 |
| Pyr → | $v = c$ | 0.4533 | ---- | 5 | *fum2* |
| Pyr → | $v = c$ | 0.4138 | ---- | 5 | C24 |
| Pyr → | $v = c$ | 0.5137 | ---- | 30 | Col-0 |
| Pyr → | $v = c$ | 0.5333 | ---- | 30 | *fum2* |
| Pyr → | $v = c$ | 0.5172 | ---- | 30 | C24 |
| TP → | $v = c$ | 0.1394 | ---- | 20 | Col-0 |
| TP → | $v = c$ | 0.1245 | ---- | 20 | *fum2* |
| TP → | $v = c$ | 0.1159 | ---- | 20 | C24 |

| | | | | | |
|---|---|---|---|---|---|
| TP → | $v=c$ | 0.1050 | ---- | 5 | Col-0 |
| TP → | $v=c$ | 0.09678 | ---- | 5 | *fum2* |
| TP → | $v=c$ | 0.1057 | ---- | 5 | C24 |
| TP → | $v=c$ | 0.1362 | ---- | 30 | Col-0 |
| TP → | $v=c$ | 0.1218 | ---- | 30 | *fum2* |
| TP → | $v=c$ | 0.1300 | ---- | 30 | C24 |
